# Metabolic engineering of anthocyanin pathway in *Artemisia annua* cells does not exclude the expression of artemisinin pathway

**DOI:** 10.1101/2023.01.05.522882

**Authors:** Rika Judd, Yilun Dong, Xiaoyan Sun, Yue Zhu, Mingzhuo Li, De-Yu Xie

## Abstract

*Artemisia annua* is the only medicinal crop to produce artemisinin for the treatment of malignant malaria. Unfortunately, hundreds of thousands of people still lose their life every year due to the lack of sufficient artemisinin. Artemisinin is considered to result from the spontaneous autoxidation of dihydroartemisinic acid in the presence of reactive oxygen species (ROS) in an oxidative condition of glandular trichomes (GTs),; however, whether increasing antioxidative compounds can inhibit artemisinin biosynthesis in plant cells is unknown. Anthocyanins are potent antioxidants that can remove ROS in plant cells. To date, no anthocyanins have been structurally elucidated from *A. annua*. In this study, our goals were to engineer anthocyanins in *A. annua* cells and to understand the artemisinin biosynthesis in anthocyanin-producing cells. Arabidopsis *PAP1* (*AtPAP1*) was used to engineer four types of transgenic anthocyanin-producing *A. annua* (TAPA1 to 4) cells. Three wild type cell types were developed as controls. TAPA1 cells produced the highest contents of total anthocyanins. LC-MS analysis detected 17 anthocyanin or anthocyanidin compounds. Crystallization, LC/MS/MS and NMR analyses identified cyanidin, pelargonidin, one cyanin, and one pelargonin. An integrative analysis characterized that four types of TAPA cells expressed the artemisinin pathway and TAPA1 cells produced the highest artemisinin and artemisinic acid. The contents of arteannuin B were similar in seven cell types. These data showed that the engineering of anthocyanins does not eliminate the biosynthesis of artemisinin in cells. These data allow us to propose a new hypothesis that enzymes catalyze the formation of artemisinin from DHAA in non-GT cells. These findings show a new platform to increase artemisinin production via non-GT cells of *A. annua*.

## Introduction

Artemisinin is an effective antimalarial sesquiterpene lactone molecule. Artemisinin-combination therapy (ACT) forms the first front treatment of malaria (WHO 2010, 2018). In addition, this compound was reported to have anti-cancer (Lai et al. 2013) and anti-diabetic activities (Li et al. 2017). Despite the effectiveness of ACT in treating malaria, this treatable disease caused the loss of about 627,000 lives in 2020 and between 558,000-896,000 from 2020-2021 (WHO 2021). This is because the production of artemisinin cannot meet the high demand of ACT for therapy. This problem is caused by its low and variable production in *Artemisia annua*, the only natural resource (Shretta and Yadav 2012; Paddon et al. 2013; Xie 2016). In addition, glandular trichomes (GTs), a 10-cell extrusion structure of the epidermis of leaves and flowers, have been considered the only cells of *A. annua* to be able to produce this compound (Duke et al. 1994a; Xie et al. 2016). This dogma has not been challenged until our recent report proved the biosynthesis of artemisinin non-GT cells (Judd et al. 2019). Our finding allows exploring new cell types to improve the production of artemisinin *in planta*. To date, the biosynthetic pathway of artemisinin has gained extensive investigations. It is derived from the mevalonate pathway located in the cytosol, and the past studies have characterized seven pathway genes encoding proteins that catalyze the formation of artemisinin (Fig. 1). In brief, amorpha-4,11-diene synthase (ADS) catalyzes the conversion of farnesyl diphosphate (FPP) to amorpha-4,11-diene (Mercke et al. 2000). A cytochrome p450 monooxygenase (CYP71AV1) and its redox partner cytochrome p450 reductase (CPR1) as well as a cytochrome b enzyme convert amorphadiene to artemisinic alcohol (Teoh et al. 2006; Paddon et al. 2013), which is converted to artemisinic aldehyde by an alcohol dehydrogenase (ADH1) (Paddon et al. 2013). An aldehyde dehydrogenase (ALDH1) dehydrogenates artemisinic aldehyde to form artemisinic acid (Teoh et al. 2009). A double bond reductase 2 enzyme (DBR2) hydrogenates artemisinic acid to form dihydroartemisinic aldehyde (Zhang et al. 2008), which is further dehydrogenated to produce dihydroartemisinic acid (DHAA) by ALDH1 (Teoh et al. 2009). A spontaneous photooxidation of DHAA is considered the last step to form artemisinin (Wallaart et al. 1999; Sy and Brown 2002a). In addition to *CPR1*, another cytochrome p450 reductase, *CPR2*, was identified from our sequence assembly and was highly expressed in floral tissues positively correlating with artemisinin accumulation in plants (Ma et al. 2015).

**Fig. 1.**
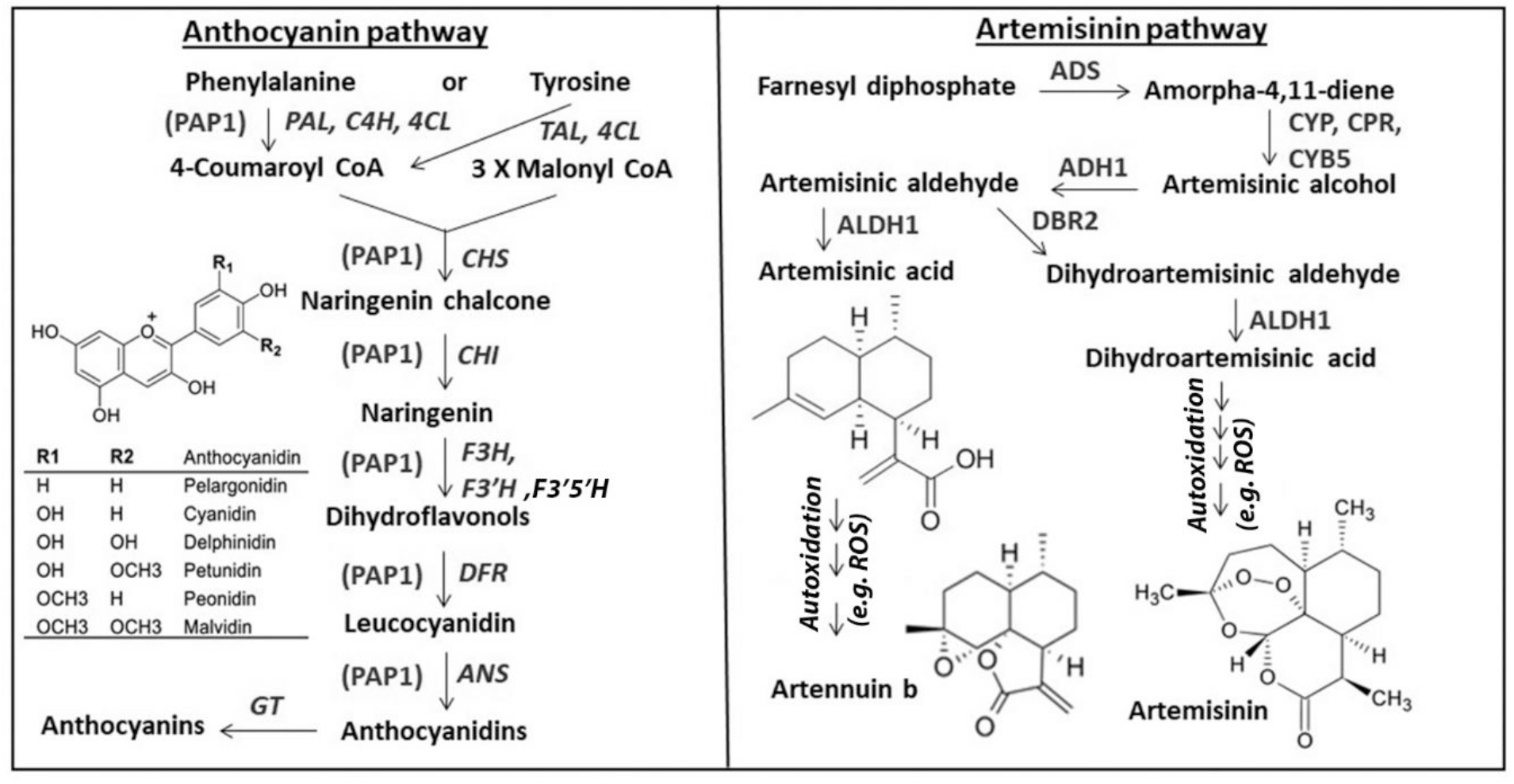
The Biosynthetic Pathways of Anthocyanins and Artemisinin in the cytosol. **A, a**nthocyanin biosynthesis begins with phenylalanine (main start) or tyrosine (minor start). Genes abbreviations, phenylalanine ammonia lyase (PAL); cinnamate 4-hydroxylase (C4H); 4-coumaroyl CoA ligase (4CL); chalcone synthase (CHS); chalcone isomerase (CHI); flavanone 3-hydroxylase (F3H); flavanoid 3’-hydroxylase (F3’H); dihdroflavonol reductase (DFR); anthocyanidin synthase (ANS); 3-glucosyltransferase (3-GT). B, the artemisinin pathway starts with farnesyl diphosphate. Question marks mean that steps open for studies. Gene abbreviation, amorpha-4,11-diene synthase (ADS); Cytochrome p450 monooxygenase 71AV1 (CYP); Cytochrome p450 reductase (CPR); Cytochrome B_5_ (CYB5); alcohol dehydrogenase (ADH1); double-bond reductase 2 (DBR2); aldehyde dehydrogenase (ALDH1). ROS: reactive oxygen species.

Anthocyanins are a group of pink/red/purple/blue flavonoid pigments with strong antioxidative activities that protect plants from stress-induced damages (Kano et al. 2005; Yuzuak and Xie 2022; Shi and Xie 2014). To date, the biosynthesis of anthocyanins has gained intensive studies in biochemistry, genetics, metabolic engineering, and applications (Shi and Xie 2014; Holton and Cornish 1995; Li et al. 2022; Zhang et al. 2014). All pathway genes and most of regulatory genes have been characterized from both model and crop plants (Shi and Xie 2014; Khusnutdinov et al. 2021). The biosynthetic pathway of anthocyanins mainly starts with phenylalanine in most plants, while it can begin with tyrosine in certain plants (Fig. 1). Pathway genes include entry (*PAL, TAL, C4H*, and *4CL*), early (*CHS, CHI, F3H, F3’H, F3’5’H*), and late pathway (*DFR, ANS*, and *GT*) genes (Fig. 1). This pathway leads to the biosynthesis of three main common groups of anthocyanidins, pelargonidin, cyanidin, and delphinidin, from which plants produce three common anthocyanin groups, pelargonin, cyanin, and delphinin. In addition to pathway genes, most regulatory genes have been characterized from different plant species. *Production of Anthocyanin Pigment 1* (*PAP1*) that is one of master regulatory genes encodes a R2R3-MYB transcription factor (Borevitz et al. 2000; Shi and Xie 2011; Li et al. 2022). In Arabidopsis, it can enhance the expression of entry and early pathway genes and activate later pathway genes (Shi and Xie 2010; Shi and Xie 2011). Therefore, *PAP1* has been selected to engineer crops for value-added traits. In roses, the *AtPAP1* overexpression increased volatile compounds and create new scent that was reported to be distinguishable by honeybees and humans (Zvi et al. 2012). The *AtPAP1* overexpression decreased alpha and beta acids in hops (Gatica-Arias et al. 2012). *AtPAP1* and Arabidopsis *transparent testa 8* were introduced into commercial tobacco plants to decrease tobacco alkaloids and carcinogens (Li et al. 2022). AtPAP1 was successfully used to engineer the anthocyanidin pathway of proanthocyanidins for animal nutrition (Xie and Dixon 2005).

To our knowledge, no anthocyanin structures have been reported from *A. annua* although more than 600 compounds including multiple flavonoids were structurally elucidated from different tissues (Brown 2010). The lack of knowledge of *A. annua* anthocyanins likely result from no production in leaves and flowers, especially GTs and other trichomes. Only a few studies reported absorbance analysis of anthocyanins from juvenile seedlings or plants or a mutant calli (Liu et al. 2017; Pandey and Pandey-Rai 2014b; Kayani et al. 2021). In addition, a few studies reported the cloning of entry and early pathway genes, including *A. annua* PAL (*AaPAL*) (Zhang et al. 2015), chalcone isomerase (*AaCHI*) (Graham et al. 2010), flavanone 3-hydroxylase (*AaF3H*) (Xiong et al. 2016). Dihydrokaempferol is hydroxylated to dihydroquercetin by flavonoid 3’-hydroxylase (F3’H). Late pathway genes have not been reported in biochemistry and genetics.

In this study, we aimed to test a hypothesis and complete two goals. The hypothesis is that *A. annua* cells can biosynthesize artemisinin in an antioxidative subcellular condition. The first goal was to understand the types of anthocyanins that *A. annua* cells could biosynthesize. Given that the formation of artemisinin requires the presence of oxidative condition rich in reactive oxygen species (ROS), the second goal was to understand whether *A. annua* cells rich in antioxidative anthocyanins could exclude the formation of artemisinin. We took advantage of AtPAP1, R2R3-MYB transformation factor (Xie et al. 2006; Shi and Xie 2011; Li et al. 2022), to engineer anthocyanins in *A. annua* cells. We developed four types of transgenic anthocyanin-producing *A. annua* (TAPA) cells, detected 17 anthocyanin or anthocyanidin compounds, from which four were identified to be cyanidin and pelargonidin, one cyanin, and one pelargonin, and characterized the biosynthesis of artemisinin in anthocyanin-producing TAPA cells.

## Results

### Development of four types of transgenic anthocyanin-producing A. annua (TAPA) calli

Agrobacterium mediated transformation and selection of transgenic cells obtained four types of TAPA calli, TAPA1, 2, 3, and 4 (Fig. 2B). TAPA1 calli were composed of soft and dark reddish cells; TAPA2 calli consisted of hard and dark reddish cells but were slightly lighter than TAPA1 cells; TAPA3 calli were formed by soft and lightly reddish cells; and TAPA4 calli were composed of hard reddish/greenish cells. Three types of wild type (WT) calli were also developed as controls. WT1 calli were hard and greenish; WT2 calli were soft and lightly yellowish/greenish; and WT3 calli were soft and lightly pale yellowish. Results of qRT-PCR analysis showed the expression of *AtPAP1* in all TAPA types of cells. The expression level of *AtPAP1* was the highest in TAPA1 cells followed by TAPA2, and TAPA3 and TAPA4 cells, the last two with a similar level (Fig. 2 D). Further estimation of total anthocyanins showed that the content was the highest in TAPA1 cells, followed by TAPA2, TAPA3, and then TAPA4. Although the expression level of *AtPAP1* was similar in TAPA3 and TAPA4 cells, the content of total anthocyanins was higher in TAPA3 than in TAPA4. By contrast, the expression of *AtPAP1* was not detected in three WT calli (Fig. 2D). These data indicated that the production of anthocyanins resulted from the overexpression of *AtPAP1*.

**Fig. 2.**
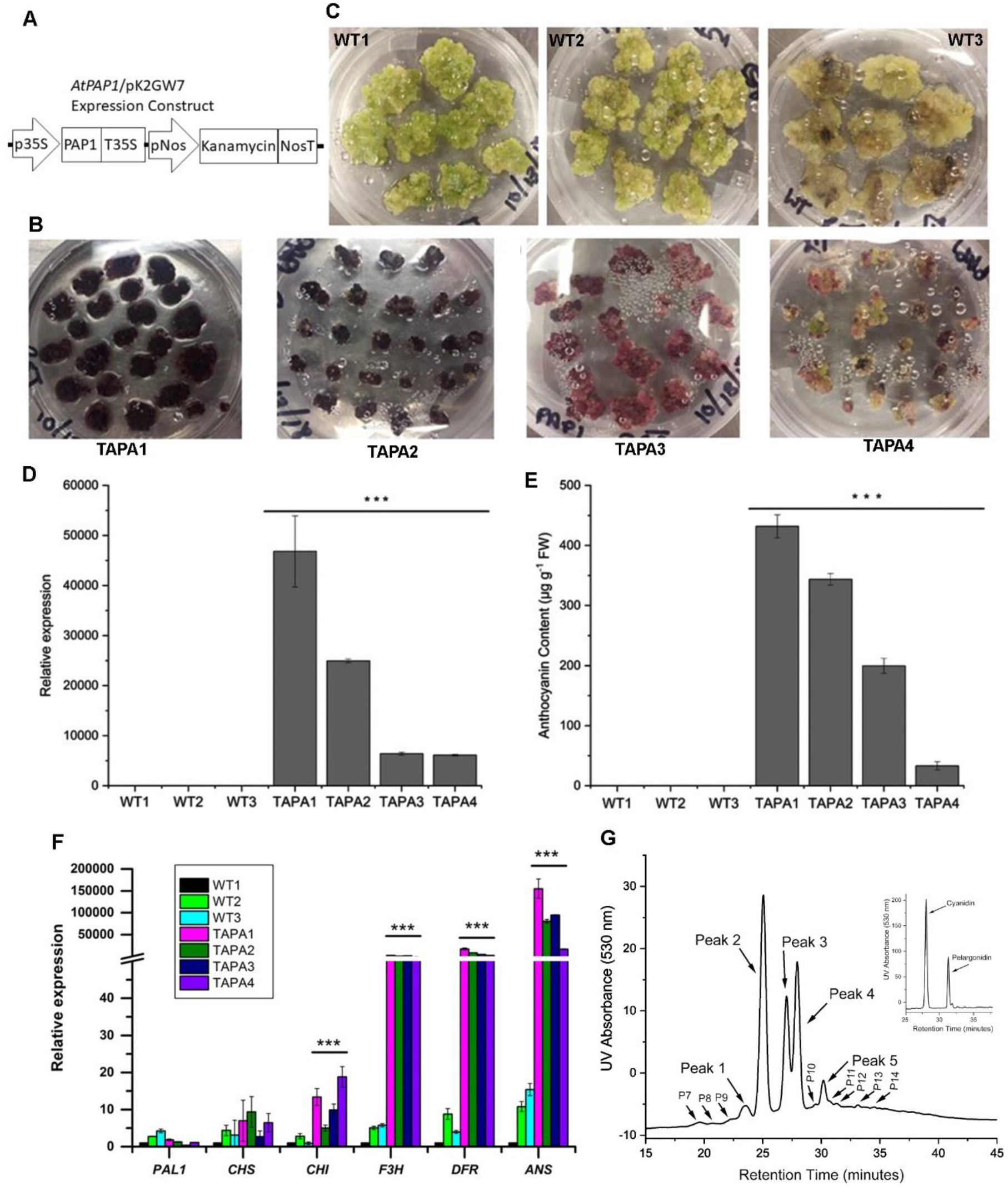
Development of the transgenic anthocyanin producing *A. annua* cells (TAPA). **A**, an AtPAP1/pK2GW7 vector was constructed from pK2GW7 to overexpress *AtPAP1* in *A. annua* cells. TAPA calli were developed after transformation of leaves and selection by antibiotics. WT calli were developed from leaves as control. All calli were subcultured three months. B, four types of red TAPA calli were developed from leaves of *A. annua*, TAPA1, TAPA2, TAPA3, and TAPA4. C, three types of wild type (WT) calli were developed from leaves, WT1, WT2, and WT3. D, Relative expression levels of the *AtPAP1* transgene in TAPA calli were estimated with qRT-PCR. The results of qRT-PCR did not show its expression in wild type (WT) calli. E, four types of TAPA calli produced different anthocyanin contents, while WT calli did not produce anthocyanins. Student’s T-test was performed to evaluate statistical significance (p-value less than 0.05). The three asterisks indicate that the TAPA cells are significantly different from all WT samples. F, relative expression of key pathway genes involved in the anthocyanin biosynthesis in TAPA calli. AtPAP1 up-regulated *CHI, F3H, DFR* and *ANS* in TAPA calli compared with wild type calli. A statistical evaluation was completed with student’s T-test. The p-value less than 0.05 indicates the significant difference. The three asterisks indicate that the TAPA cells are significantly different from all WT samples. G, Anthocyanin profiles in extracts of TATA1 cells were detected by HPLC recorded at 530 nm. Five main (Peak 1-5) and more than nine peaks (P6-14 and more) were detected at 530 nm. The insert chromatogram shows cyanidin and pelargonidin produced from the hydrolysis of TAPA1 anthocyanins.

### Characterization of anthocyanin biosynthesis in TAPA cells

To characterize the anthocyanin pathway engineered in TAPA cells, we completed qRT-PCR to analyze the expression of *PAL1, CHS, CHI, F3H, DFR*, and *ANS*, which were associated one entry step, three early steps, and two late steps (Fig. 1). Results showed the high expression of *F3H, DFR*, and *ANS* in all TAPA cells but only a background level of these three genes in wild type (WT) cells (Fig. 2 F). The expression levels of *CHI* were significantly higher in TAPA cells than in WT cells. The expression levels of *CHS* were slightly higher in TAPA cells than in WT cells. The expression level of *PAL1* was not significantly increased in TAPA cells compared to WT cells. These data indicate that AtPAP1 activates later pathway gene expression and increases early pathway gene expression leading to the activation of the anthocyanin biosynthesis in *A. annua* cells.

HPLC-qTOF-MS/MS analysis was completed to characterize anthocyanin profiles in TAPA cells. The UV spectrum recorded at 530 nm detected 13 anthocyanin peaks (corresponding to 17 anthocyanin or anthocyanidin molecules described below) from the extracts of TAPA1 cells (Fig. 2 G). Peak #1, 2, 3, 4, and 5 were main ones. Three other TAPA cell types also produced all of these anthocyanin peaks, although the peak sizes were apparently reduced (Fig. S1). This datum supported the results of total anthocyanin contents described above (Fig. 2 E). Hydrolysis of extracts coupled with HPLC assays revealed cyanidin and pelargonidin, two common types of anthocyanin chromophores, from the extracts of the four types of TAPA cells (Fig. 2G and Fig. S1). This result indicates that the anthocyanins engineered in TAPA cells are composed of cyanin and pelargonin. HPLC-qTOF-MS/MS analysis was further completed to characterize the [m/z]^-^ values of these peaks. In addition, fragmentations were created for each peak. The resulting data showed [m/z]^-^ values of peak# 2, 3, 4, 5, 6, and 7, but could not obtain unambiguous [m/z]^-^ values for other peaks (Table 1). The m/z]^-^ values of peak# 3, 6, and 7 were 535, 447, and 431. Based on the feature of fragmentations, peak#3 and 6 were cyanin-like anthocyanins featured with a fragment 285 corresponding to cyanidin, while peak #7 was a pelargonin-like anthocyanin characterized with a fragment 269 belonging to pelargonidin. It was interesting that two or three primary [m/z]^-^ values were identified from peak#2, 4, and 5. The peak#2 was characterized with 551 and 533 [m/z]^-^ values, from which an apparent fragment 285 was generated to correspond to cyanidin by collision induced dissociation (CID) of MS/MS, thus, they were proposed to be cyanin-like anthocyanins. The peak #4 included three primary 641, 575, and 285 [m/z]^-^ values, from the first and second of which an apparent 285 fragment was also generated by CID, thus, they were proposed to be cyanin like anthocyanins. The 285 [m/z]^-^ value was directly associated with cyanidin. The peak #5 included two primary 533 and 269 [m/z]^-^ values, from the first of which an apparent fragment 269 was generated by CID, thus, it corresponded to pelargonidin and was proposed to be a pelargonin like anthocyanin. The 269 [m/z]^-^ value corresponded to pelargonidin. These data indicate that the engineered anthocyanins are composed of cyanin and pelargonin.

**Table 1.** [m/z]^-^ characterization of anthocyanins from extracts of TAPA Cells by HPLC-MS/MS analysis.

| Peak (P) # | m/z [M-2H]<sup>-</sup> | Retention Time | Major MS/MS Fragments | Predicted Structure |
| --- | --- | --- | --- | --- |
| 1 | Unidentified | 23.9 | Unidentified | Unidentified |
| 2 | 551 [cy]<sup>-</sup> | 25.250 | 267, **285**, 307, 329, 371, 381, 397, 471 | Cyanin like anthocyanin |
| 2 | 533 [cy]<sup>-</sup> | 25.318 | 284, **285**, 297, 363, 422, 447, 463, 490 | Cyanin malonyl-glucoside like anthocyanin |
| 3 | 535 [pel]<sup>-</sup> | 26.972 | **269**, 287, 291, 329, 355, 371, 397, 449, 491 | Pelargonin like anthocyanin |
| 4 | 641 [cy]<sup>-</sup> | 28.109 | 191, 207, 284, **285**, 307, 345, 367, 389, 429, 471, 500, 561 | Cyanin like anthocyanin |
| 4 | 575 [cy]<sup>-</sup> | 28.101 | 284, **285**, 327, 444, 471, 531 | Cyanin like anthocyanin |
| 4 | 285 [cy]<sup>-</sup> | 27.999 | **285** | Cyanidin |
| 5 | 533 [pel]<sup>-</sup> | 30.238 | 259, **269**, 282, 287, 310, 313, 325, 340, 355, 369, 440 | Pelargonin like anthocyanin |
| 5 | 269 [pel]<sup>-</sup> | 31.33 | 269 | Pelargonidin |
| 6 | 447.09 [cy]<sup>-</sup> | 19.8 | 125, 213, **284**, **285**, 349 | Cyanidin-3-O-glycoside |
| 7 | 431.12 [pel]<sup>-</sup> | 21.7 | 107, 147, 199, 241, **269**, 419 | Pelargonidin-3-O-glycoside |
| 8 | Unidentified | 21.8 | Unidentified | Unidentified |
| 9 | Unidentified | 29.5 | Unidentified | Unidentified |
| 10 | Unidentified | 31.8 | Unidentified | Unidentified |
| 11 | Unidentified | 32.1 | Unidentified | Unidentified |
| 12 | Unidentified | 32.9 | Unidentified | Unidentified |
| 13 | Unidentified | 34.6 | Unidentified | Unidentified |

### Isolation of two anthocyanins and two anthocyanidins

A scale-up culture was carried out to isolate anthocyanins for structural elucidation. Solvent fractionation and three types of columns were used to isolate individual components (Fig. 3). The first separation on a silica column obtained six main fractions (Fra. A, B, C, D, E, and F), which were further used for separation on Sephadex LH-20 columns. In this experiment, Fra. B was further selected for purification. After the use of three types of LH-20 columns, four fractions, Fra. B314-1, 2, 3, and 4 were obtained for crystallization. After Fra. B314-1, 3, and 4 were collected and their liquid volumes were reduced, crystals were formed from the remaining solution and collected for structural analysis. The crystal formed from Fra. B314-1 was labelled as B314-12. Fra. B314-2 was further separated with a XDB-C18 reverse column on HPLC. After the removal of elution solvent, we obtained about 4 mg purified B314-22 powder.

**Fig. 3.**
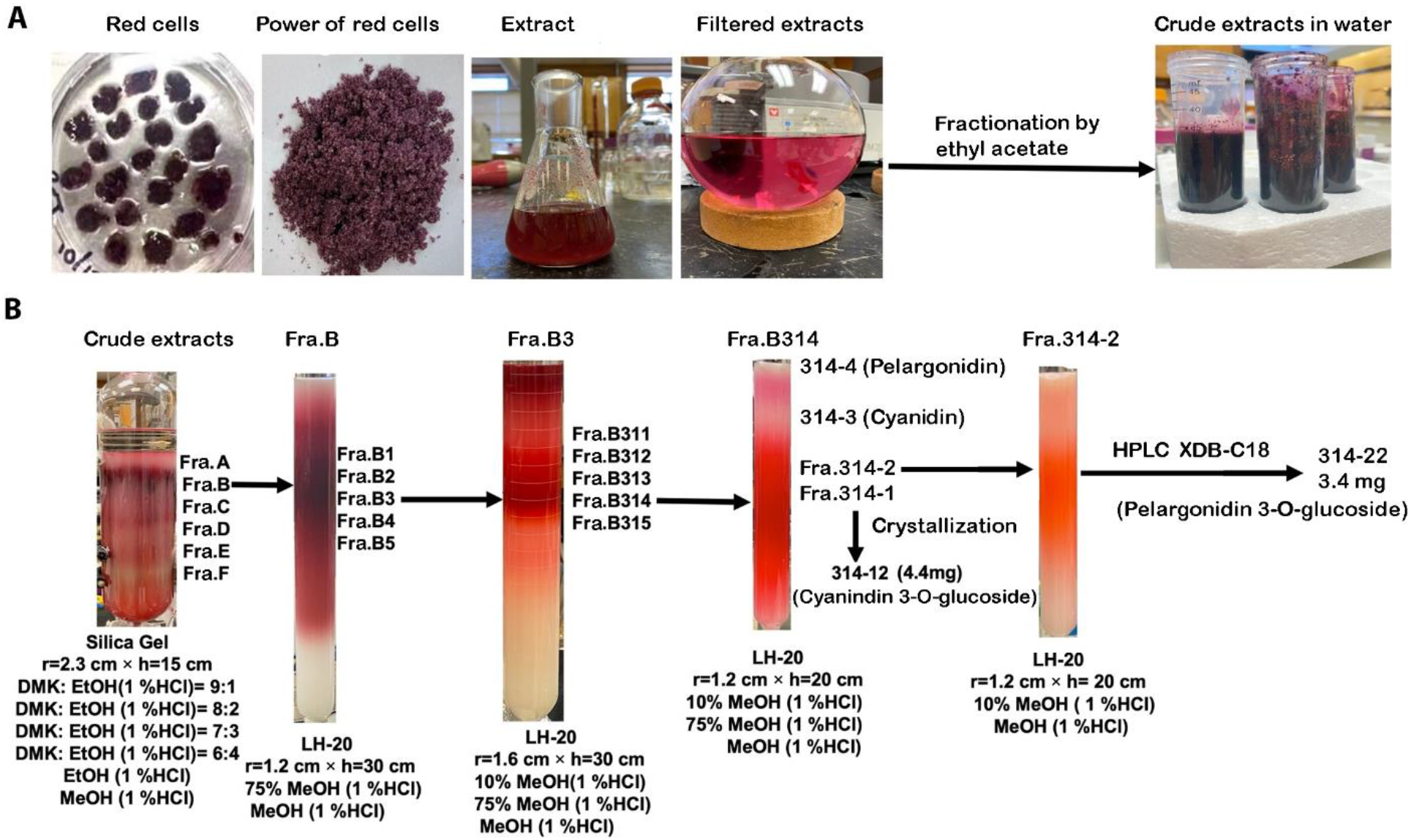
Purification steps of two anthocyanidins and two anthocyanins from extracts of TAPA1 cells. A, red calli cultured on solid medium were used to extract crude anthocyanins, which were re-extracted with ethyl acetate to remove non-polar compounds to obtain anthocyanins soluble in water. B, A silica gel column, LH-20 Sephadex gel columns, and a C18 reverse phase silica column were used to obtain different fractions that led to the isolation of four fractions, fraction 314-3 (cyanidin), fraction 314-4 (pelargonidin), fraction 314-12 (cyanidin 3-O-glucoside), and fraction 314-22 (pelargonidin 3-O-glucoside).

HPLC-MS/MS analysis was further carried out to characterize the molecular weights of these four isolates. The results showed that Fra. B314-12 and B314-22 were composed of single peak and their [m/z]^-^ values were 447 and 431 (Fig. 4A and B). The feature fragments from CID of Fra. B314-12 and B314-22 were 284 or 285 and 269. This datum indicated that Fra. B314-12 and B314-22 were a cyanin and a pelargonin molecule, respectively. Furthermore, HPLC-MS analysis and comparison with standard samples identified that B314-3 and B314-4 crystals were cyanidin and pelargonidin, respectively (Fig. 4C and D).

**Fig. 4.**
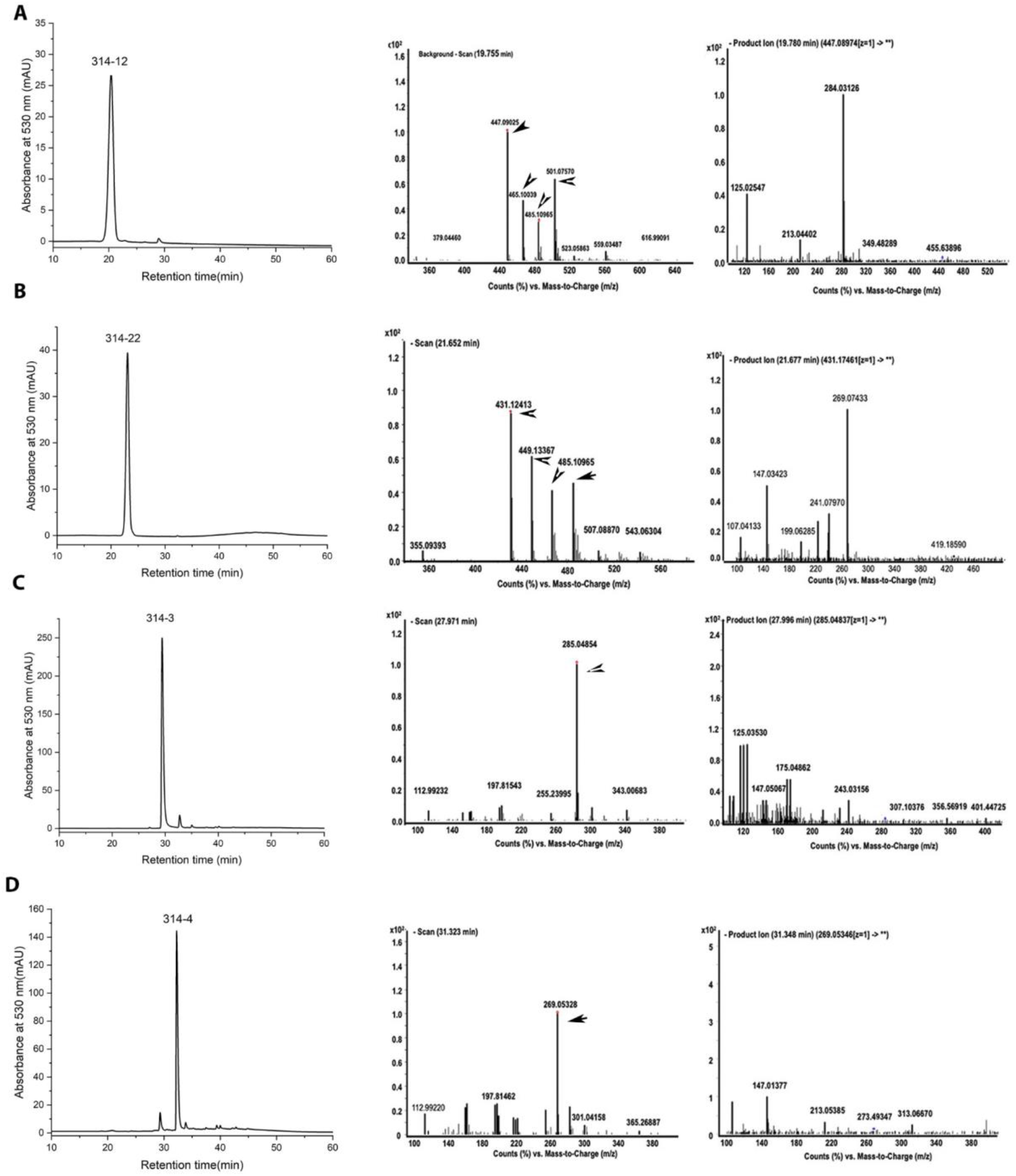
HPLC-MS/MS analysis of four fractions. HPLC profiles, ESI features, and MS/MS fragment patterns of 314-12 (A), 314-22 (B), 314-3 (C) and 314-4 (D) fractions. HPLC profiles were recorded at both 280 nm and 530 nm. The [m/z]^-^ values of 314-12 (A), 314-22 (B), 314-3 (C) and 314-4 (D) are 447, 431, 285, and 269, respectively.

### Structural elucidation of Fra. B314-12 and B314-22 by NMR

1D1H, 1H-1H correlation spectroscopy (COSY) and heteronuclear multiple bond correlation experiments (HMBC) NMR tests of Fra. B314-11 and -22 were completed on a Bruker NEO 700MHz spectrometer equipped with a TCI Cryo-Probe. The spectra of both 1H, 1H-1H, 13C, and 1H-13C were recorded (Figs. 5 and 6, Figs. S2-S5). These analyses allowed obtaining the chemical shift assignments of 1H and 13C of the two isolates. Based on these analyses, two compounds were identified to consist of 21 carbons. Fra. B314-12 and -22 had 21 and 19 hydrogens. All 13Cs and 1Hs were assigned (Table 2). Based on the assignment, Fra. B314-12 and 22 were identified to be cyanidin-3-O-glycoside and pelargonidin-3-O-glycoside (Fig. 5 A and B).

**Table 2:**
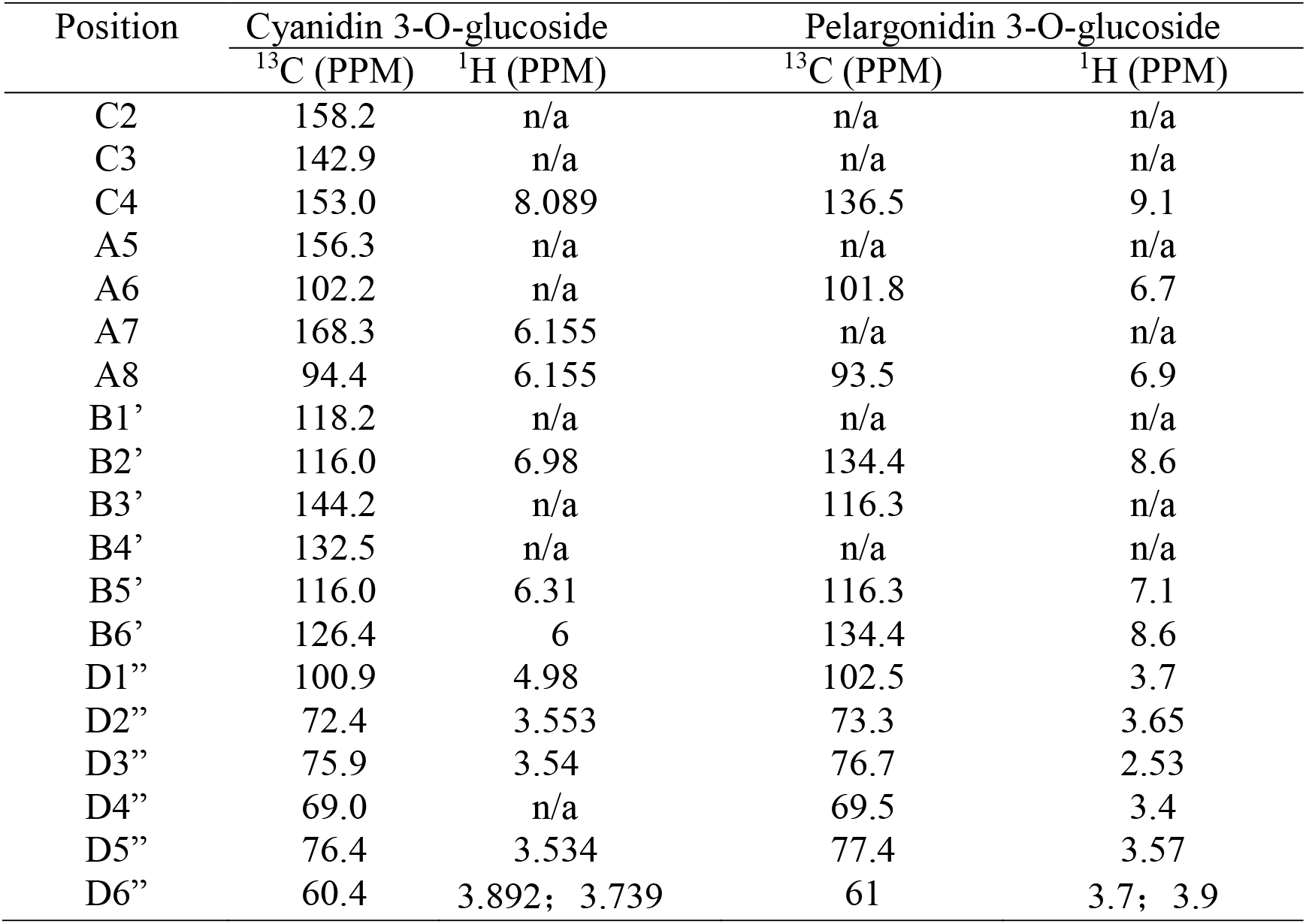
^13^C and ^1^H NMR assignments of cyanidin 3-O-glucoside and pelargonidin 3-O-glucoside

**Fig. 5.**
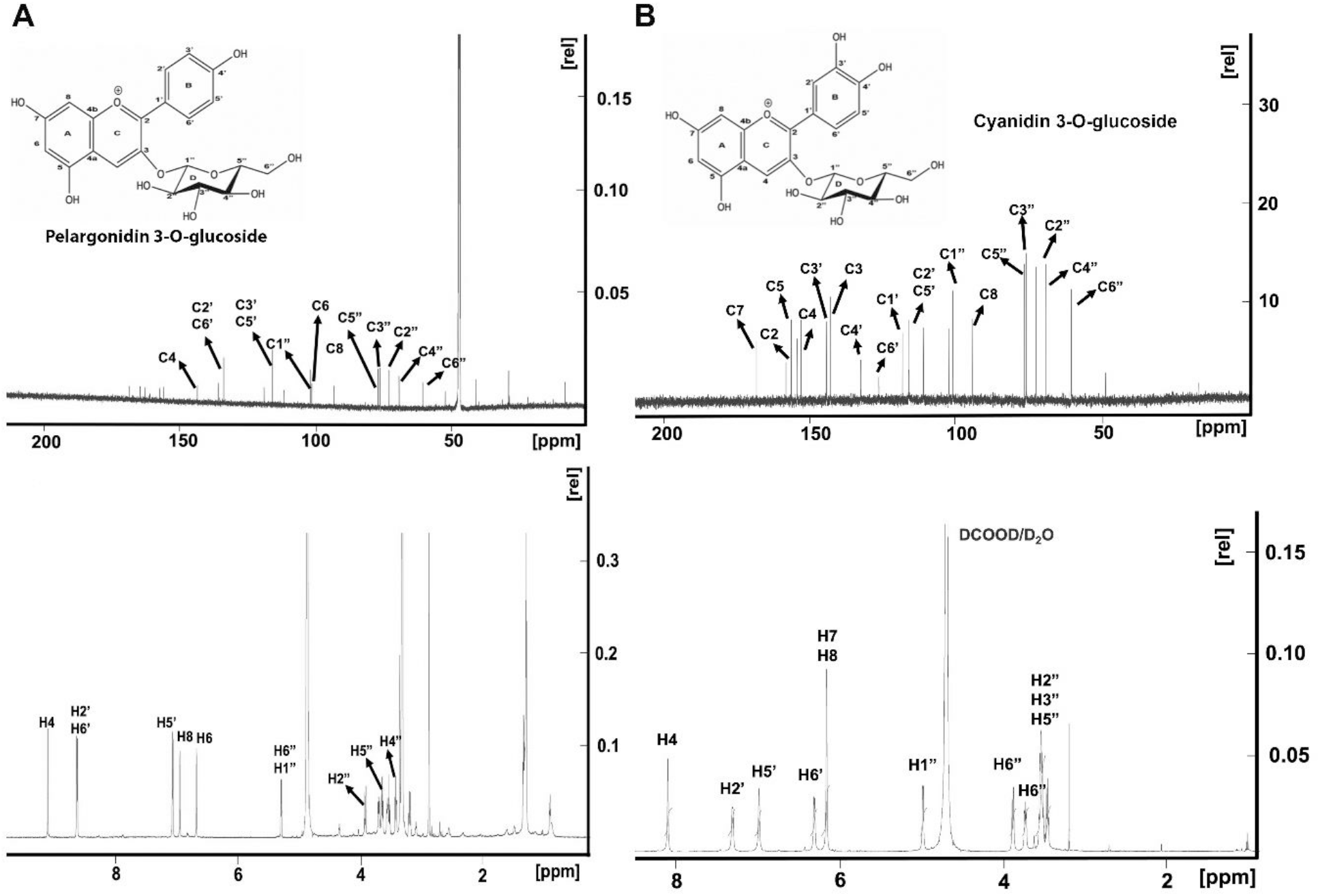
^13^C and ^1^H NMR spectra. A, two plots show features of ^13^C (upper) and ^1^H (bottom) NMR spectra of 314-22 and identify it to be pelargonidin 3-O-glucoside. B, two plots show features of ^13^C (upper) and ^1^H NMR (bottom) spectra of 314-12 and identify it to be cyanidin 3-O-glucoside.

**Fig. 6.**
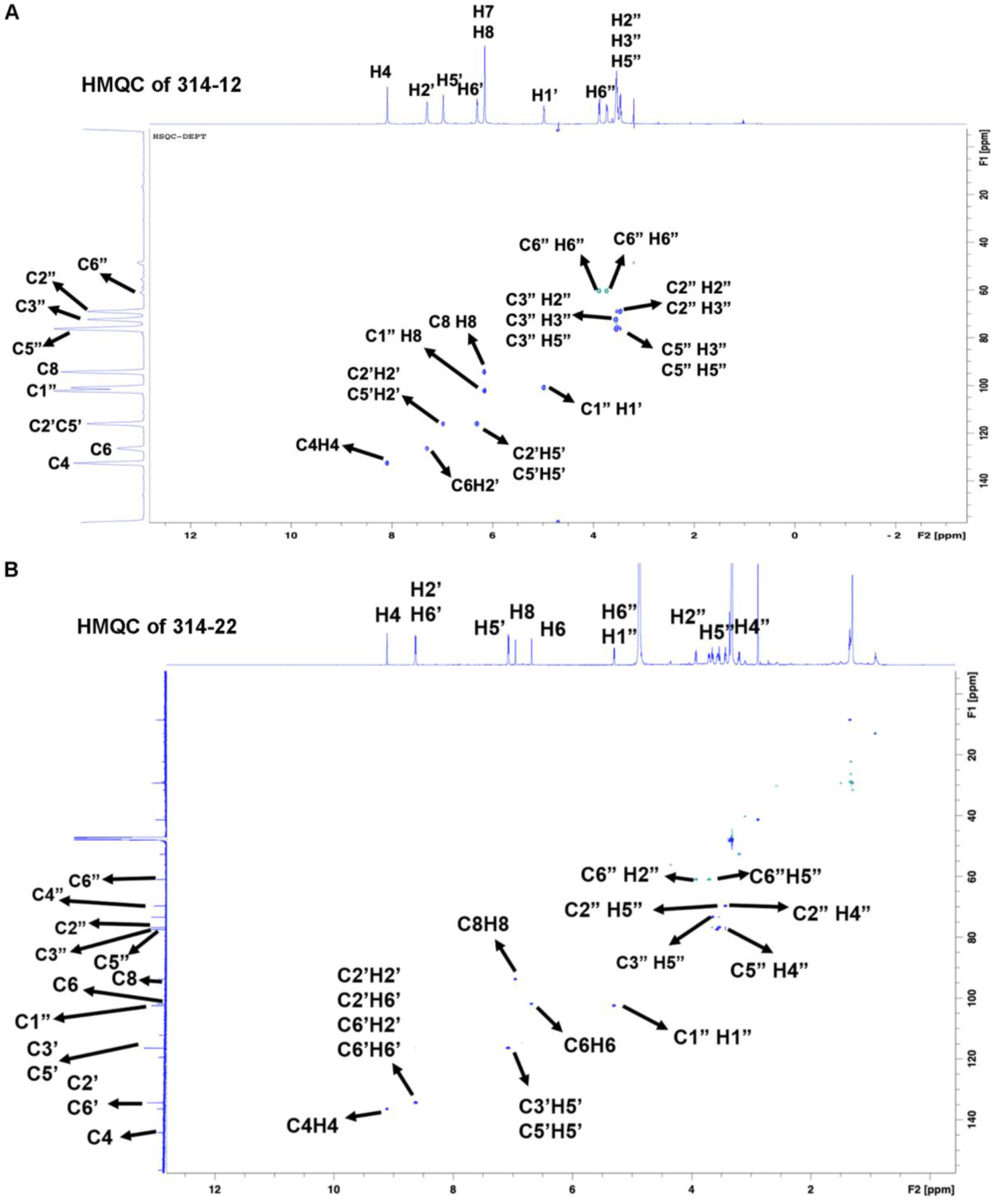
Heteronuclear single-quantum correlation spectroscopy (HSQC) of NMR of fractions 314-12 and 314-22. A-B, two plots show features of HSQC NMR spectra of fractions 314-12 (A) and 314-22 (B), which are identified to be cyanidin 3-O-glucoside and pelargonidin 3-O-glucoside.

### Characterization of the artemisinin pathway in TAPA cells

We performed gene expression and metabolic profiling to understand artemisinin biosynthesis in TAPA cells. Results from qRT-PCR analyses showed that four TAPA cell types and three WT cell types expressed six known pathway genes, *ADS, CYP71AV1, CRP1, ADH1, DBR2*, and *ALDH1* (Fig. 7 A). The expression of genes in WT cell types supported our previous gene expression data in calli (Judd et al. 2019). Although the expression levels of these gene varied in four TAPA cell types and three WT cell types, the results indicated that the expression levels of *CYP71AV1* was upregulated in all TAPA cell types, while the expression levels of *CPR1* were downregulated in all TAPA cell types (Figure 7A). The expression levels of other genes were associated with each cell types. Compared with WT1 cells, the expression levels of *ADS, ADH1, DBR2*, and *ALDH1* were either increased or decreased in four TAPA cell types and two other WT cell types. *CPR2* was previously reported to potentially associate with the artemisinin biosynthesis in flowers (Ma et al. 2015). This gene was expressed in three WT cell types but was hardly detected in all four TAPA cell types. To demonstrate the activity of the artemisinin pathway, HPLC-MS/MS analysis was performed to identify artemisinin, artemisinic acid, and arteannuin B. The resulting data showed that four TAPA and three WT cell types produced these three metabolites (Fig. 7 B-D, Fig. S6-S8). The contents of artemisinin and artemisinic acid were higher in TAPA1 cells than in other cell types (Fig. 7 B and C). By contrast, the contents of artemisinin were lower in TAPA2, TAPA3, and TAPA4 cells than in three WT cell types. The contents of arteannuin B were similar in seven cell types (Fig. 7D). All data indicate that four TAPA cell types engineered with anthocyanin production do not exclude the biosynthesis of artemisinin, artemisinic acid, and arteannuin B.

**Fig. 7.**
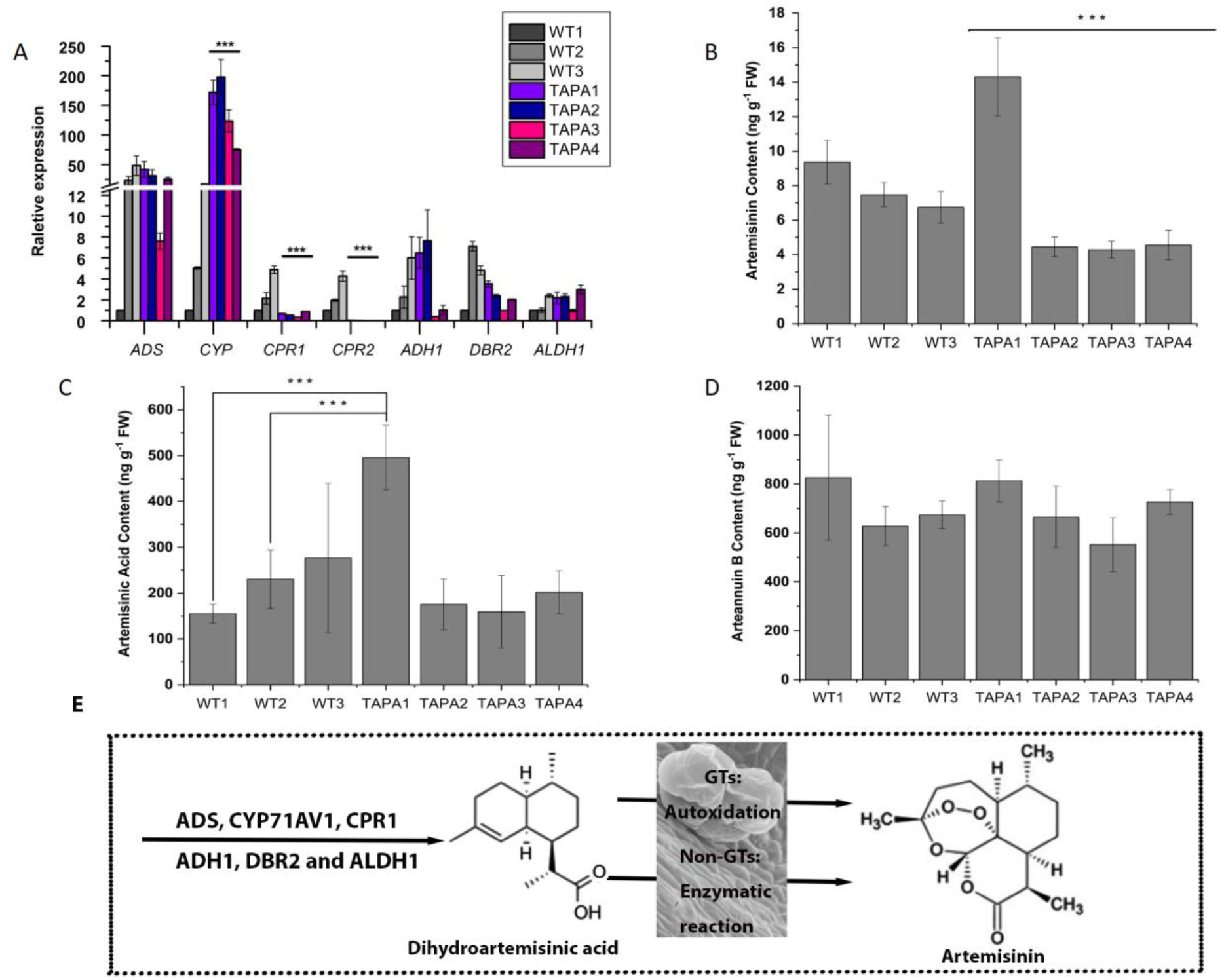
Artemisinin Biosynthesis in TAPA Cells. **A**, qRT-PCR data show the relative expression levels of six known artemisinin pathway genes. The resulting data indicated that both red and wild type cells expressed six artemisinin pathway genes, the level of *CYP71 (CYP)* was upregulated in all TAPA cells, and the expression of *CPR2* was detected in wild type cells but not in red cells (A). **B**, the artemisinin content in TAPA1 was the highest, while those in TAPA2, TAPA3, and TAPA4 were lower than in three types of wild type cells (WT1, 2, and 3). The three asterisks over all TAPA cells indicate that they are significantly different from the three WT samples. **C**, the content of artemisinic acid was higher in TAPA1 than in TAPA2, TAPA3, TAPA4, WT1, and WT2 cells, while was not significantly different from that in WT3 cells. Three asterisks over TAPA1 and WT1 and WT2 indicate significantly difference. D, the contents of arteannuin b content were similar in all cell types. E, a hypothesis is that enzymes are responsible for the conversion of dihydroartemisinic acid to artemisinin in non-glandular trichome cells. Students’ T-test was performed to evaluate the statistical significance between two samples e.g. TAPA1 and one of other cell types. A p-value less than 0.05 indicates the significance.

## Discussion

The first goal of this study was to understand what types of anthocyanins could be produced in *A. annua*. Of over 600 plant secondary metabolites structurally elucidated from *A. annua* (Brown 2010), about 49 are flavonoids. To date, however, no anthocyanin structures have been elucidated from *A. annua* by either crystallization or NMR although a few studies reported either spectrometer-based absorbance estimation or the tissue’s pigmentation of anthocyanins (Hong et al. 2009; Srivastava and Sangwan 2012; Pandey and Pandey-Rai 2014a, b; Kumar et al. 2016; Liu et al. 2017). Therefore, it is interesting to understand anthocyanin types that *A. annua* can biosynthesize. We have observed that anthocyanin pigmentation mainly occurs in hypocotyls of juvenile seedlings with limited biomass; accordingly, we used *AtPAP1* to engineer cells to produce anthocyanins. Four types of red cells overexpressing *AtPAP1* (Fig. 2) were developed to characterize anthocyanin biosynthesis. HPLC-MS/MS analysis detected 17 anthocyanins or anthocyanidins formed in engineered TAPA cells (Table 1). Of them, four were purified via column chromatography under this report (Fig. 3) and then identified to be two anthocyanidins, pelargonidin and cyanidin, and two known anthocyanins, cyanin-3-O-glycosides and pelargonidin-3-O-glycoside (Figs. 4-6 and Figs. S1-S5). Although the structures of 13 other anthocyanins remained unknown, hydrolysis of total anthocyanins released cyanidin and pelargonidin without delphinidin (Fig. 2 G and Fig. S1). This datum indicates that the unknown 13 anthocyanins (Table 1) are composed of either cyanin or pelargonin. In addition, pathway gene expression analysis showed that the overexpression of *AtPAP1* increased the expression of *A. annua (Aa) CHI* and activated *AaF3H, AaDFR*, and *AaANS* (Fig. 2 F). This datum characterizes the biosynthetic pathway of anthocyanins from phenylalanine to anthocyanidins (Fig. 1) in *A. annua*. Taken together, the engineering of TAPA cells for the first time provides insight into the structural chromophore of anthocyanins and their biosynthesis in *A. annua*.

The second goal underlying this study was to understand whether an antioxidative condition created by anthocyanins excluded the formation artemisinin and arteannuin B, the products of autoxidation. To date, although the last step of artemisinin formation remains for further investigation, the only hypothesis accepted globally is that the spontaneous autoxidation of dihydroartemisinic acid (DHAA) produces artemisinin (Fig. 1) (Sy and Brown 2002a, b; Czechowski et al. 2016; Brown 2010; Paddon et al. 2013; Czechowski et al. 2022). Both chemical conversion and plant treatments and senescence have provided evidence to support this hypothesis. A four-step autoxidation was developed to convert DHAA to artemisinin, the reaction of the Delta (4,5)-double bond of DHAA with oxygen, Hock cleavage, oxygenation, and cyclization (Sy and Brown 2002a). Moreover, this autoxidation technology formed a main platform in the development of synthetic biology of artemisinin from artemisinic acid that was engineered from yeast (Ro et al. 2006; Paddon et al. 2013). The hypothesis that the autoxidation-based formation of artemisinin occurs *in planta* has resulted from the observations of stress-associated conditions. The treatment of *A. annua* plants with a high light intensity increased reactive oxygen species (ROS) that significantly enhanced the production of artemisinin. A burst of singlet oxygen was reported to associate with the increase of artemisinin in senescent leaves (Yang et al. 2010). The treatment of UV-B that increased ROS was also reported to increase artemisinin production in plantlets. A seasonal increase of artemisinin in leaves was proposed to result from a ROS-based conversion of DHAA (Wallaart et al. 2000). All these reports are solely based on the accepted theory that the biosynthesis of artemisinin only occurs in glandular trichomes (GT) consisting of 10 secretary cells, the outgrowth of epidermis of leaves and flowers (Duke et al. 1994b; Czechowski et al. 2016; Zhang et al. 2008; Xiao et al. 2016; Covello et al. 2007; Xie et al. 2016). Based on the reports, it can be proposed that a reductive condition rich in antioxidants in the cytosol of GTs may inhibit or exclude the biosynthesis of artemisinin. In addition to GTs, our recent findings showed that non-GT cells expressed the entire artemisinin pathway to produce artemisinin (Judd et al. 2019). Another recent study reported that the production of most artemisinin mainly resulted from non-GTs (Liao et al. 2022). If the biosynthesis of artemisinin in non-GT cells also results from autoxidation, it is interesting to understand if a creation of antioxidative condition can inhibit or exclude the formation of this antimalarial compound. Anthocyanins are strong antioxidative pigments (Kano et al. 2005; Mohammad et al. 2021; Bishayee et al. 2010). Especially, the biosynthesis of anthocyanins is highly induced by stress conditions, such as UV-radiation (Hatier and Gould 2008; Merzlyak et al. 2008; Rowan et al. 2009; Shi and Xie 2014; Yuzuak and Xie 2022) and anthocyanins have strong scavenging capacity to remove ROS (Fan et al. 2023; Pang et al. 2023; Qu et al. 2018; Zhang et al. 2016). Based on previous reports, our hypothesis is if an antioxidative condition can inhibit the biosynthesis of artemisinin, the engineering of anthocyanins likely excludes the formation of artemisinin in non-GT cells. To test this hypothesis, we used *AtPAP1* to develop four types of anthocyanin-producing cells (Fig. 2. B). LC-MS analysis showed that although these cells produced anthocyanins (0.1 mg-0.8 mg/g FW, equivalent to 0.1-0.8% (dry weight) (Fig. 2 E), all these cells expressed the pathway genes of artemisinin and produced artemisinin, artemisinic acid, and arteannuin B (Fig. 7). It was apparently noted that the contents of these three compounds depended upon cell types. TAPA1 cells produced the highest contents of anthocyanins, artemisinin, and artemisinic acid. The contents of artemisinin, artemisinic acid, and arteannuin B were no significantly different among three types of WT cells, TAPA2, TAPA3, and TAPA4. These data suggest that the engineering of high anthocyanin production did not deactivate the biosynthesis of artemisinin, artemisinic acid, and arteannuin B. In another words, an antioxidative cellular condition does not exclude the artemisinin pathway and its activities.

Based on these findings, we propose that non-GT cells and GTs have different mechanisms associated with the last steps of artemisinin biosynthesis (Fig. 7 E). In GTs, as discussed above, the autoxidation of DHAA forms artemisinin. The extrusion of GTs may obtain sufficient oxygen and have sufficient amount of singlet oxygen, which lead to oxidation of DHAA to form artemisinin. In non-GTs, we propose that enzymatic reactions are responsible for the conversion of DHAA to artemisinin. The evidence is that TAPA cells with high production of anthocyanins do not exclude the biosynthetic pathway of artemisinin. Especially, the artemisinin content of TAPA1 cells with the highest contents of anthocyanins (Fig. 2) was higher than that of WT cells (Fig. 7 B). Given that antioxidative anthocyanins are biosynthesized in the cytosol and then transported to the central vacuole (Shi and Xie 2014), our TAPA cell data indicate that the biosynthesis of artemisinin most likely occurs in the absence of an oxidative condition. Which part of explants did TAPA cells originate from, parenchymal, spongy cell, palisade, epidermal, or hypodermal cells? Our previous studies reported that when the overexpression of *AtPAP1* was used to engineer anthocyanins, microscopic observations determined that epidermal and hypodermal cells, parenchymal cells in vascular bundles, and linear trichomes were the biosynthetic locations (Shi and Xie 2010; Xie et al. 2006; Broeckling et al. 2012). Based on these reports, we hypothesized that TAPA cells likely originated from epidermal cells, hypodermal cells, or/and parenchymal cells in vascular bundles. The reason is that these cells, such as epidermal cells, are active locations for induction of cell division to form calli (Xie and Hong 2001). By contrast, TAPA cells were unlikely from GTs. The reason is that GTs are completely differentiated cells and are unlikely to be transformed by *Agrobacterium tumefaciens*-based transformation. Furthermore, our recent MS imaging data indicated that artemisinin-producing non-GT cells were most likely from epidermis (Judd et al. 2019), suggesting that TAPA cells were likely associated with epidermal origin. A recent report quantified artemisinin in GTs alone, non-GT cells, and GTs/non-GT mixtures (Liao et al. 2022). The resulting data (shown in their supporting Figure 1 by Liao et al 2022) reported that non-GTs accounted for more than 90% artemisinin in leaf tissues including GTs/non-GT cells. Because of the biomass limitation of GTs in plants, this observation shows a promise for the improvement of artemisinin production in the futures. It can be anticipated that elucidating the last steps of artemisinin in non-GT cells will enhance metabolic engineering of artemisinin in *A. annua* for high production.

## Materials and methods

### Induction of AtPAP1 transgenic cells

We previously used gateway cloning to clone the full-length cDNA of *AtPAP1* into the pK2GW7 plant expression vector to obtain a AtPAP1/pK2GW7 vector (Fig. 2A), which was introduced into the *Agrobacterium tumefaciens* GV3101 strain for genetic transformation (He et al. 2017). In addition, pK2GW7 (empty vector) was introduced to GV3101 as a control. For transformation, the *AtPAP1*-pK2GW7/GV3101 and pK2GW7/GV3101 cultures grew on a rotary shaker to an OD_600_ of 1.556 and 1.5, under 28°C 28°C at 250 rpm, respectively. The GV3101cultures were harvested by centrifugation at 4000 rpm for 5 min. The resulting cultures were then re-suspended in 70 ml of MS liquid medium and then shaken at 250 rpm at 28°C for 1 hr of activation. Meanwhile, 30-day old sterile self-pollinated *A. annua* plants grown on MS medium in the jars were used to collect leaf explants. Leaves were cut into 1×1 cm^2^ pieces, which were co-incubated with the activated GV3101 suspensions contained in sterile petri dishes. The infected explants were dabbed on sterile filter papers to remove excessive Agrobacterium GV3101 prior to inoculate them on MS medium containing 1 mg/L BAP, 0.05 mg/L NAA, 100 μM acetosyringone, and 2 g/L of gelrite, which were contained in petri dishes. Then, all petri dishes were placed in the dark for co-cultivation. After 3 days, the co-cultivated explants were washed three times with sterile water to remove *Agrobacterium* and then placed on sterile filter papers to remove excessive water. All washed explants were inoculated on a selection medium consisting of MS medium supplemented with 1 mg/L BAP, 0.05 mg/L NAA, 200 mg/L timentin, 40 mg/L kanamycin, and 2 g/L of gelrite contained petri dishes, which were placed on tissue culture shelves under a 16/8 (light/dark) photoperiod (light intensity: 50 μmol/min, m^2^) and 25°C. After three weeks, red cells were induced from a number of explants and formed small apparent calli, which were transferred to freshly prepared selection medium and then subcultured every two weeks for one month. In addition, calli were induced from explants infected by empty vector control transformation. Wild type (WT) calli as controls were induced from explants without transformation.

Four steps were completed to select a medium for propagation of TAPA calli with high production of anthocyanins. First, all calli were transferred onto medium I consisting of basal MS medium supplemented with 0.5 BAP and 0.5 NAA, and then were subcultured every two weeks for three months. Second, all calli were cultured on medium II consisting of basal MS medium supplemented with 0.25 mg/L 2,4-dichlorophenoxyacetic acid (2,4-D) and 0.1 mg/L kinetin and subcultured every two weeks for one month. Third, the concentration of 2,4-D in medium II was increased to 0.5 mg/L without a modification of other components to obtain medium III. All calli were transferred onto medium III and subcultured every two weeks for 4 months. Finally, all calli were transferred onto medium IV consisting of basal MS medium supplemented with 1 mg/L NAA, 0.1 mg/L kinetin, and 1 mg/L of ascorbic acid. After 4 months of subculture on medium IV, four types of red calli transformed by *AtPAP1*-pK2GW7/GV3101 were developed and named as transgenic <u>a</u>nthocyanin-<u>p</u>roducing *<u>A</u>rtemisia annua* (TAPA) cells, which had different levels of red pigmentation (Fig. 2B). In addition, three types of WT calli were obtained as controls (Figure 2C). Vector control transgenic calli were also developed as controls.

### Semi-quantitative and real-time quantitative RT-PCR assays

Different TAPA and WT cell lines were used for gene expression analysis. Fresh samples were ground into a fine powder in liquid nitrogen and stored in a -80°C freezer. As described in our previous report, total RNA was extracted from 100 mg powdered sample with the IBI isolate (IBI Scientific, USA) kit by following the manufacturer’s instructions, treated with DNase to remove genomic DNA, and then stored in a -80°C freezer.

The first strand cDNAs were synthesized from all DNA-free RNA samples with a high capacity cDNA reverse transcription kit (Applied Biosystems, USA) by following the manufacturer’s instructions, and then used for real time RT-PCR assays. Gene specific primer pairs were designed for *AtPAP1, AaPAL1, AaCHS, AaCHI, AaF3H, AaF3’H, AaDFR*, and *AaANS* for amplification and the thermal cycles of semi-quantitative RT-PCRs were designed for each gene (Supplemental Table 1). qRT-PCR was completed with the iTaq-Universal SYBR Green Supermix by following manufacturer’s instructions (Bio-Rad, USA). In addition to analyzing the expression levels of the 8 anthocyanin pathway genes, the relative expression levels of six artemisinin pathway genes, *AaADS, AaCYP, AaCPR1, AaADH1, AaDBR2*, and *AaALDH1*, were analyzed in all samples. *AaCPR2*, a hypothetical gene involved in the biosynthesis of artemisinin, was also analyzed. Gene specific primer pairs for *AaADS, AaCYP, AaCPR1, AaADH1, AaDBR2*, and *AaALDH1* have been previously reported (Judd et al. 2019). The primer pair for *AaCPR2* is included in Supplemental Table 1. *Beta-ACTIN* was used as an internal control for normalization. The steps of qRT-PCR assay were the same as described previously (Judd et al. 2019). In brief, DNA-free RNA samples were diluted 3 times prior to amplification. The thermal cycle consisted of 50°C for 10 min, 95°C for 10min, and 45 cycles of 95°C for 15 secs and 60°C for 1 min. Three technical replicates were performed for each RNA sample. The ΔΔC_t_ algorithm was used to calculate relative expression levels. The expression levels of genes in TAPA cells were then normalized with those in WT1 cells.

### Extraction of anthocyanins, measurement, and hydrolysis

The extraction method of anthocyanins with 0.5% HCl in methanol and the measurement of total contents were the same as described previously (He et al. 2017). In brief, 100 mg (fresh weight, FW) of ground powder from each biological sample was used to extract anthocyanins. Three biological samples were extracted for each cell line. The final volume of each sample extract was 2 ml. The absorbent values of 1 ml anthocyanin extracts were measured at 530 nm on a spectrophotometer. Three technical replicates were performed for each biological sample. A standard curve was developed with cyanidin chloride in a concentration range from 6.25 to 100 ng/ml). Based on the standard curve, the total anthocyanin content in each biological sample was estimated as cyanidin equivalent. Anthocyanin extracts were hydrolyzed to identify anthocyanidins. The methods of hydrolysis and sample preparation were the same as previously described (Shi and Xie 2011). Finally, each hydrolyzed anthocyanin sample was dissolved in 1 ml of 0.1% HCl in methanol. Both anthocyanin and anthocyanidin extracts were stored at -20°C for HPLC-qTOF-MS/MS analysis described below.

### Extraction of artemisinin and its related metabolites

Artemisinin and its related metabolites were extracted from samples. The steps of extraction and the measurement approaches were the same as previously described (Alejos-Gonzalez et al. 2011). In brief, 200 mg (FW) of ground powder of each biological sample was used to extract metabolites with ethyl acetate/acetonitrile (50:50, volume/volume). Three biological samples were extracted for each cell line. Extracts were dissolved in LC-MS grade methanol and the final volume was 400 μl. All extracts were stored in -20°C for HPLC-MS analysis as described below.

### High performance liquid chromatography-quadruple-time-of-flight tandem mass spectrometer (HPLC-qTOF-MS/MS) analysis

HPLC-qTOF-MS/MS analysis was performed on Agilent 1200 HPLC coupled with 6520 time-of-flight MS/MS (Santa Clara, CA, USA) to analyze anthocyanins. A protocol developed to annotate or identify flavanols (Yuzuak et al. 2018) was used for analysis of anthocyanin and anthocyanidin. In brief, samples were separated on an Eclipse XDB-C18 analytical column (250 mm x 4.6 mm, 5 μm, Agilent, Santa Clara, CA, USA). The mobile phase solvents were composed of 1% acetic acid in water (solvent A) (HPLC-grade acetic acid and LC-MS grade water) and 100% acetonitrile (solvent B) (LC-MS grade). To separate metabolites, the following gradient solvent system composed of ratios of solvent A to B was used, including 90:10 (0–5 min), 90:10–88:12 (5–10 min), 88:12–80:20 (10–20 min), 80:20–55:45 (20–45 min), 55:45– 50:50 (45–55min), 50:50–90:10 (55–60 min), 90:10–10:90 (60–60.01min), and 10:90–10:90 (60.01-70 min). After each run was finished, the column was washed 10 min with 10 % solvent B for equilibration. The elution rate was 0.4 ml/min. The injection volume was 10 μl. The drying gas flow was 12 l/min, and the nebulizer pressure was 50 psi. The chromatographic profiles of anthocyanins were recorded at 530 nm with a DAD detector. Metabolites were ionized with the negative mode. The mass spectra were scanned from 100 to 3000 m/z. The acquisition rate was three spectra per second. Other MS conditions included fragmentor: 150 V, skimmer: 65 V, OCT 1 RF Vpp: 750 V, and collision energy: 30 psi. Three authentic standards, cyanidin, pelargonidin, and delphinidin, were used as positive controls.

Analysis of artemisinin, artemisinic acid, and arteannuin B was also completed on Agilent 1200 HPLC coupled with 6520 time-of-flight MS/MS. A protocol was appropriately developed for identification and quantification of these compounds (Judd et al. 2019). We used this protocol to quantify the contents of these three compounds in all cell lines.

### Purification of anthocyanins from TAPA cells

The callus culture of TAPA1 cells was scaled up to obtain 500 g fresh samples (Fig. 3A). All calli were harvested and then ground into fine powder in liquid nitrogen. The resulting powder was suspended in 5 liters (L) of 1% acetic acid in 95% ethanol. The mixture were ultra-sonicated for 20 min and then filtered with 3M filter paper in an aspirator filter pump. These steps were repeated 5 times. All acetic acid-ethanol extracts were combined to obtain about 25 L, which was transferred to a rotary flask (Fig. 3A). The combined extract was evaporated to reduce the volume to about 200 ml at 65 °C with a rotary evaporator. The anthocyanin extract was concentrated in acetic acid-water. Then, 200 ml of ethyl acetate (EA) was added to the condensed extract. The resulting mixture was vortexed and centrifuged to obtain the bottom water-anthocyanin phase and the upper EA phase. The EA phase contained other phenolic compounds. These steps were repeated three times. The acetic acid water phase was moved to a clean rotary flask and evaporated to obtain solid red residue at 65 °C by a rotary evaporator. The residue was dissolved in 1% HCl methanol for HPLC analysis and purification. HPLC analysis performed on 2010EV LC/DAD instrument (Shimadzu, Japan) was to examine main anthocyanin peaks as described previously (Shi and Xie 2011). The column used for separation, mobile phase solvents, and gradient elution solvent system were the same as those used in HPLC-qTOF-MS/MS analysis described above. The chromatographs were recorded from 190 to 800 nm.

Silica gel, Sephadex LH-20, and reverse phase C18 columns were used to isolate anthocyanins (Fig. 3 B). First, 200 g silica gel (technical grade, pore size, 60 A, 230-400 mesh, particle size, 40-60um, Supelco) were loaded into a glass column (r=2.3 cm × h=15 cm), and washed with 250 ml 1% HCl in methanol, followed by equilibration with 250 ml 1% HCl in acetone/ethanol (v/v=9:1). About 13 g crude extracts dissolved in 20 ml 1% HCl in methanol were loaded onto the column. Anthocyanins were eluted with seven solvents in the order of 1: acetone/ethanol (v/v=9:1) with 1 % HCl (250 ml), 2: acetone/ethanol (v/v=8:2) with 1 % HCl (50 ml); 3: acetone/ethanol (v/v=8:2) with 1 % HCl (275 ml), 4: acetone/ethanol (v/v=8:2) with 1 % HCl and acetone/ethanol (v/v=7:3) with 1 % HCl (180 ml), 5: acetone/ethanol (v/v=6:4) with 1 % HCl, 6: ethanol with 1 % HCl (200 ml), and 7: methanol with 1 % HCl (150 ml) (Fig. 1B). The flow rate was 1 ml**/**min. The separated bands were collected to obtain multiple fractions. Each fraction was concentrated by using a rotary evaporator at 45 °C, freeze-dried, and measured for weight. The residues were dissolved in 1% HCl in methanol for HPLC analysis. Those fractions with similar anthocyanin profiles were combined to obtain six fractions, Fra. A-F. Of these, Fra. B had about 3.4 g in weight and was used for immediate purification of anthocyanins with LH-20 columns. Second, a Sephadex LH-20 chromatographic column (r=1.2 cm × h=30 cm; GE Healthcare Bio-Sciences AB) was prepared by packing powder into a glass column and washed to remove contaminants with an elution buffer consisting of 1% HCl in 75% ethanol. Fra. B was loaded onto this column, followed by elution with two types of solvents, 200 ml 75% ethanol (1% HCl) and 200 ml ethanol (1% HCl) at a flow rate of 4 ml/min. Multiple fractions were collected for HPLC analysis. Those collections with similar anthocyanin profiles were combined to obtain five fractions, B1-B5, which were freeze-dried and stored at -20 °C for further isolation. Third, a second Sephadex LH-20 chromatographic column (r=1.6 cm × h=30 cm) was prepared to isolate anthocyanins in Fra. B3. After Fra. B3 was loaded onto this column, anthocyanins were eluted with 10% methanol (1% HCl), 75% methanol (1% HCl), and then methanol (1% HCl) with a flow rate of 4 ml/min. Based on anthocyanin profiles examined by HPLC, those fractions with similar profiles were combined to obtain five fractions, Fra. B311-315, which were freeze-dried and stored at -20 °C. Forth, a third Sephadex LH-20 chromatographic column (r=1.2 cm × h=20 cm) was prepared to separate anthocyanins in Fra. B314. After this fraction was loaded onto this column, anthocyanins were eluted with 10% methanol (1% HCl), 75% methanol (1% HCl), and then methanol (1% HCl) with a flow rate of 4 ml/min to obtain four fractions, Fra. B314-1, 2, 3, and 4. Of these four, B314-1, 3, and 4 formed crystals, while B314-2 did not. Fifth, a fourth Sephadex LH-20 chromatographic column (r=1.2 cm × h=20 cm) was prepared to separate anthocyanins in Fra. B314-2. After loaded onto this column, anthocyanins were eluted with 10% methanol (1% HCl) and methanol (1% HCl) with a flow rate of 4 ml/min. One main fraction was collected. Sixth, to isolate the main anthocyanin component, this main fraction was freeze-dried and re-dissolved in 100 μl 1% HCl in methanol for isolation of anthocyanin via HPLC performed on 2010EV LC/DAD instrument. Ten-μl B314-2 sample was injected into an Eclipse XDB-C18 column (250 mm x 4.6 mm, 5 μm, Agilent, Santa Clara, CA, USA) and eluted with the same gradient solvent system described above. Fractions were collected to multiple 1.5 ml Eppendorf tubes. After multiple injections, one main peak detected at 530 nm was collected and pooled together. After the pooled collection was freeze-dried to powder, a small portion of which was re-dissolved in 1% HCl in methanol for HPLC analysis to examine its purity.

Finally, four purified anthocyanin fractions, B314-3, B314-4, B314-12, and B314-22 (Fig. 3 B), were obtained for structural elucidation via HPLC-qTOF-MS/MS described above and NMR analysis described below.

### Structure elucidation with nuclear magnetic resonance (NMR)

To elucidate the structures of Fra. B314-12 and B314-22, 1D 1H, 1H-1H correlation spectroscopy (COSY) and heteronuclear multiple bond correlation experiments (HMBC) NMR tests were performed on a Bruker NEO 700MHz spectrometer equipped with a Cryo-Probe. Fra. B314-12 (4.4 mg) was dissolved in 600μl of D_2_O and Fra. B314-22 (3.3mg) was dissolved in 600 μl of CD_3_OD. 1D 1H NMR spectra were recorded at the temperature of 240K. The 1H chemical shift and structure conformation were assigned by using 1H-1H COSY and NOESY with a mixing time of 0.8 min and the Distortionless Enhancement by Polarization Transfer-Heteronuclear Single Quantum Coherence experiment (DEPT-HSQC) and Heteronuclear Multiple Bond Correlation (HMBC) experiments performed at 240K.

### Statistical analysis

All measurements were completed with at least three biological replicates each with three technical replicates. Values were averaged from replicates and standard errors were calculated to show variations. Student t-test was used to evaluate statistical difference. P-value less than 0.05 means significant differences.

## Acknowledgement

We thank College of Agriculture and Life Sciences Dean’s Graduate Research Assistantship and U.S. Department of Education GAANN Biotechnology Fellowship, NC State University for supporting Rika Judd’s PhD study. We thank China Scholar Council for supporting Yilun Dong’s PhD Exchange Scholarship.

## Author Contributions

RJ performed genetic transformation and developed all cell types, analyzed anthocyanin pathway and artemisinin pathway, participated LC-MS analysis, and drafted the manuscript. YD performed and scaled up tissue and cell culture, extracted anthocyanins, developed isolation technology to separate anthocyanins, participated HPLC-MS/MS analysis of anthocyanins, performed NMR tests, elucidated structures of anthocyanidins and anthocyanins, and drafted the manuscript. XYS designed and tested NMR experiments and elucidated structures of anthocyanins; YZ performed LC-MS/MS experiments and data analysis; ML participated tissue culture and gene expression analysis. DYX perceived the entire project, designed and supervised all experiments, participated in data analysis, and drafted and finalized the manuscript.

## Supplementary Material

**Figure S1.**
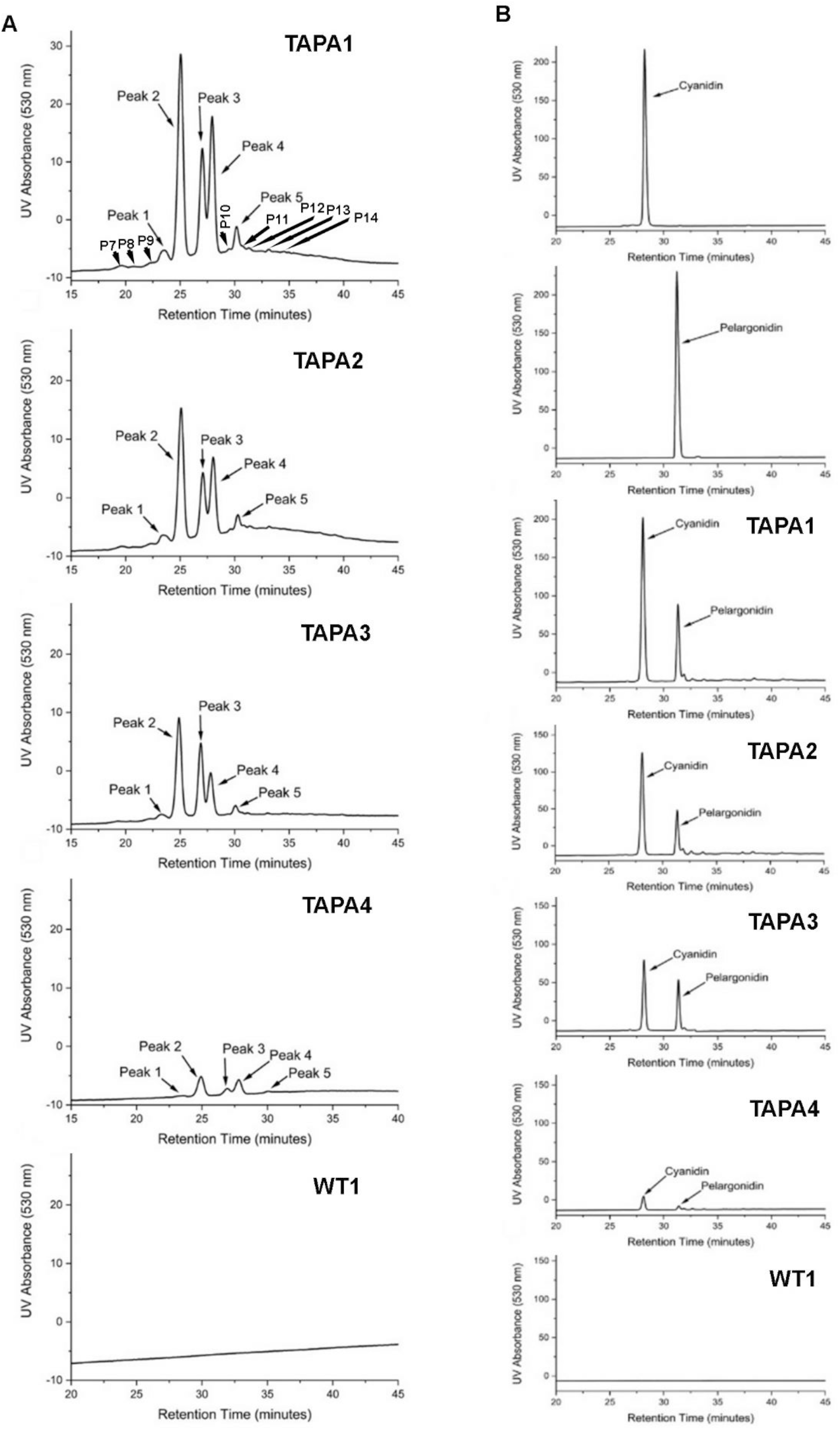
Anthocyanin profiles and main anthocyanidins in TAPA cells. A, HPLC chromatograms show different anthocyanin profiles detected extracts of TAPA1, TAPA2, TAPA3, and TAPA4 (D), while no anthocyanin peaks were detected for any extracts of WT1. No peaks were detected from extracts of WT2 and WT3 either. B, anthocyanidin profiles from the hydrolysis of anthocyanin extracts of TAPA cells. Cyanidin and pelargonidin are two standards and two were produced form the hydrolysis of anthocyanins of TAPA1, TAPA2, TAPA3, and TAPA4, while were not detected from the hydrolysis of WT1 extract.

**Fig. S2.**
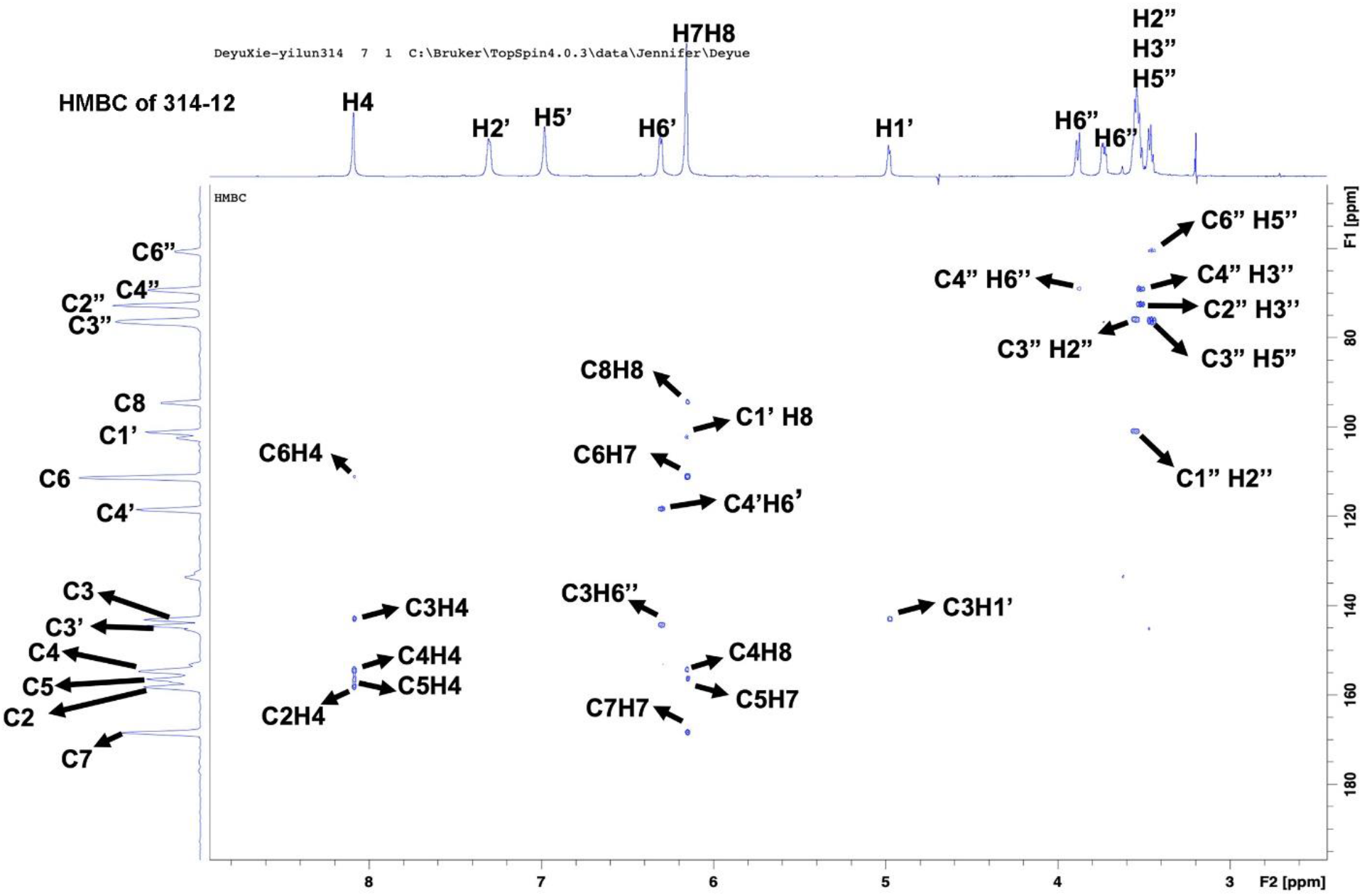
DEPT-HSQC and HMBC NMR spectrum of fraction 314-12. This spectrum indicates that fraction 314-12 is cyanidin 3-O-glucoside.

**Fig. S3.**
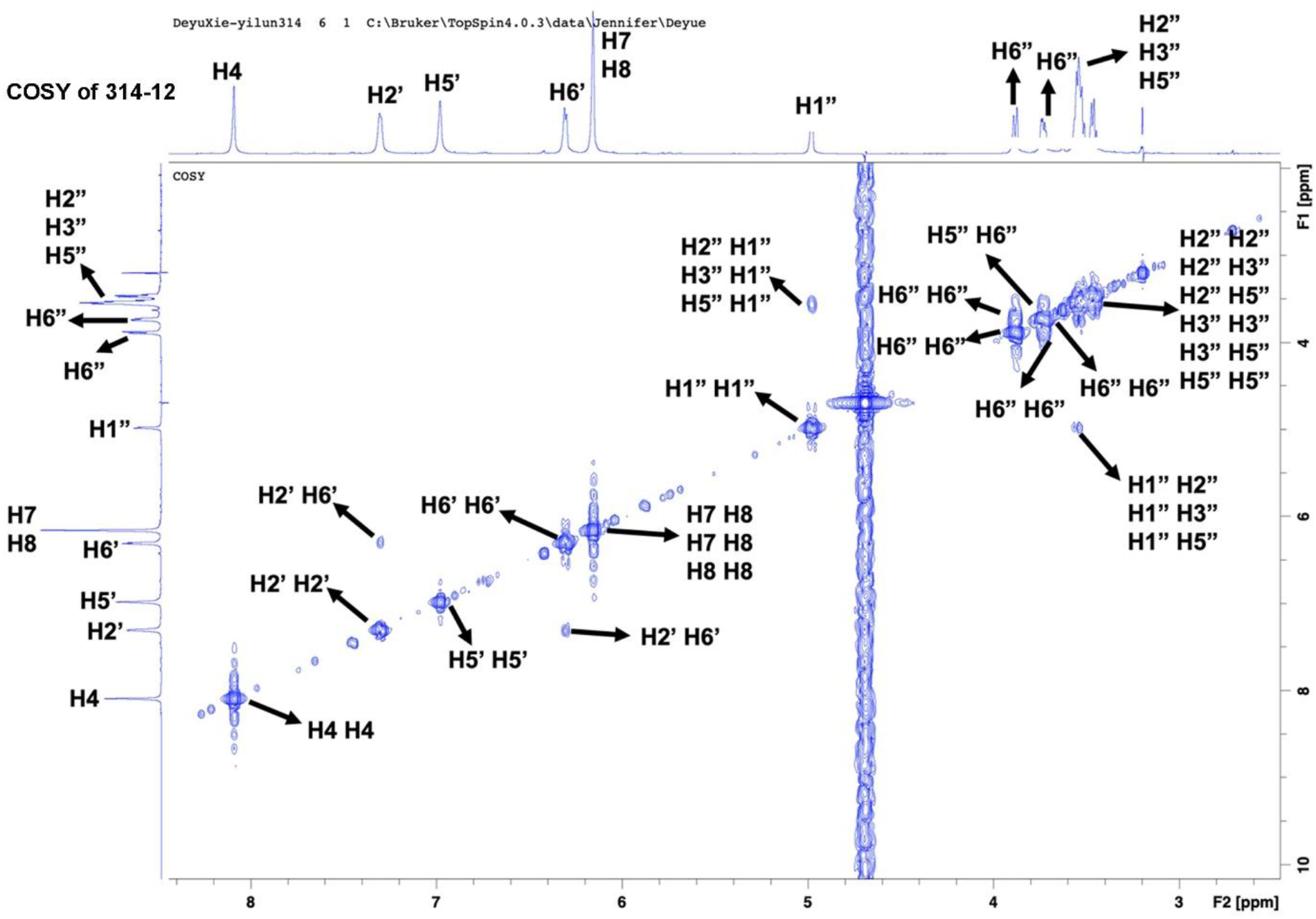
COSY NMR spectrum of fraction 314-12. This spectrum indicates that fraction 314-12 is cyanidin 3-O-glucoside.

**Fig. S4.**
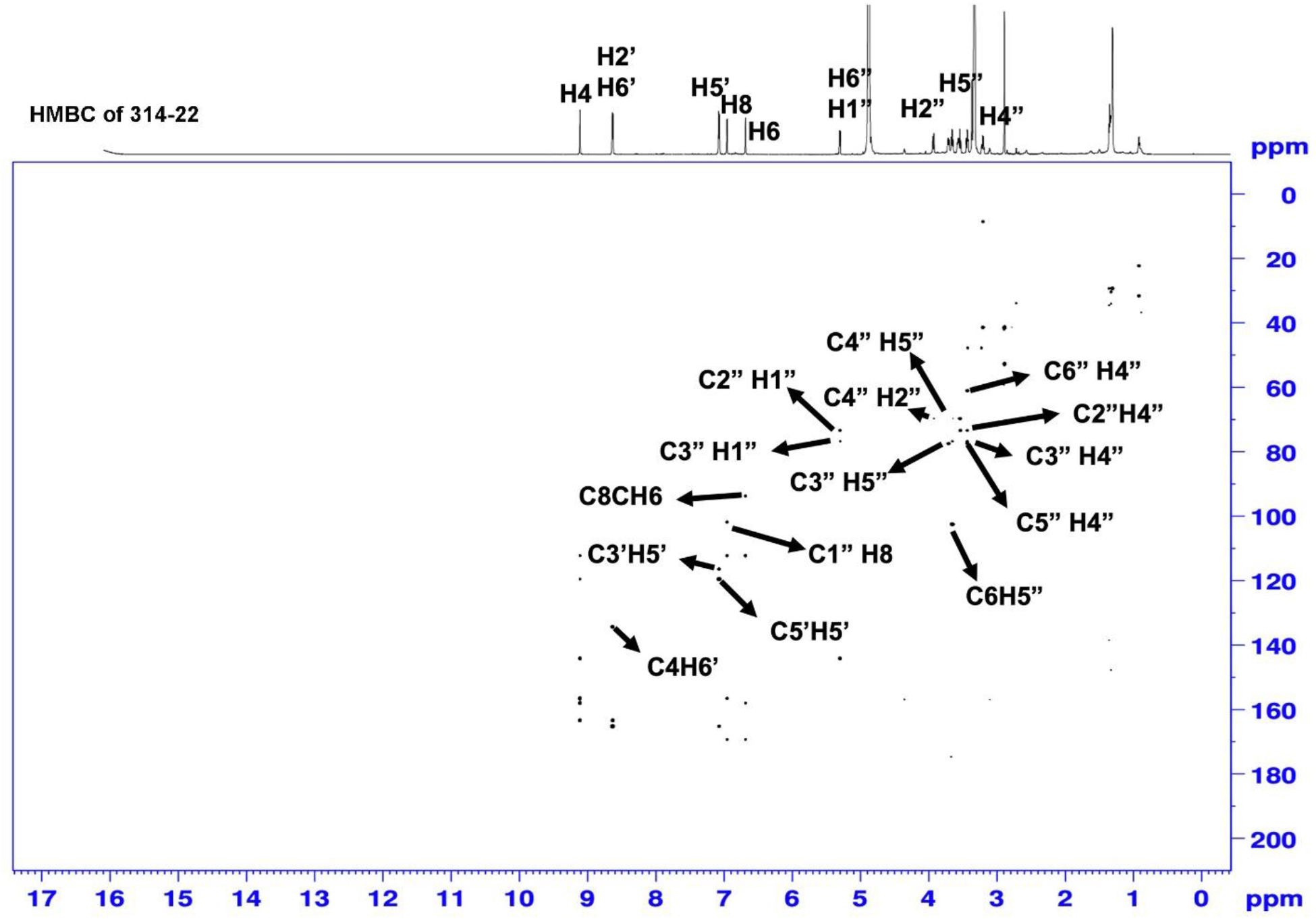
HMBC NMR spectrum of fraction 314-22. This spectrum indicates that fraction 314-22 is pelargonidin 3-O-glucoside.

**Fig. S5.**
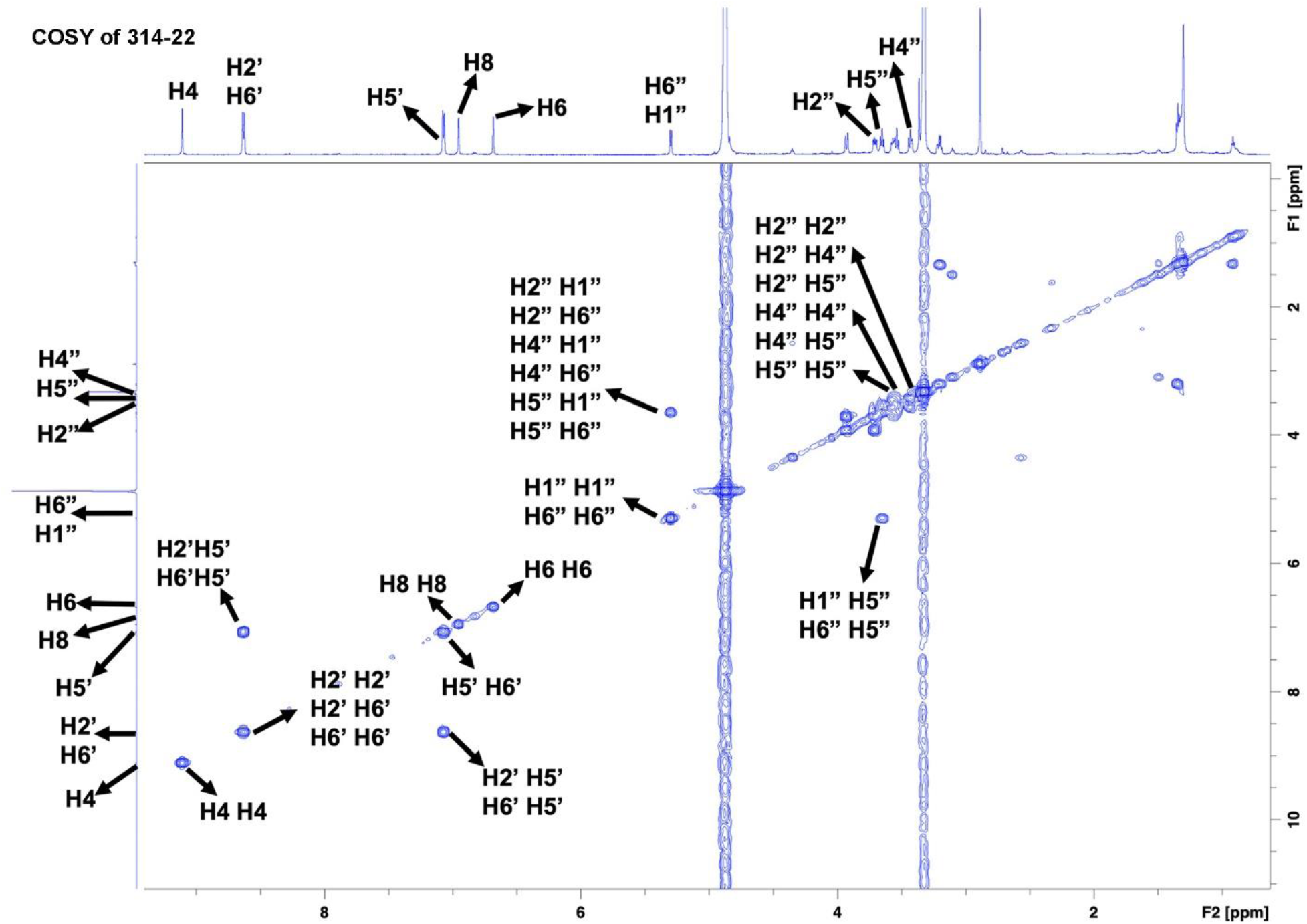
COSY NMR spectrum of fraction 314-22. This spectrum indicates that fraction 314-22 is pelargonidin 3-O-glucoside.

**Fig. S6.**
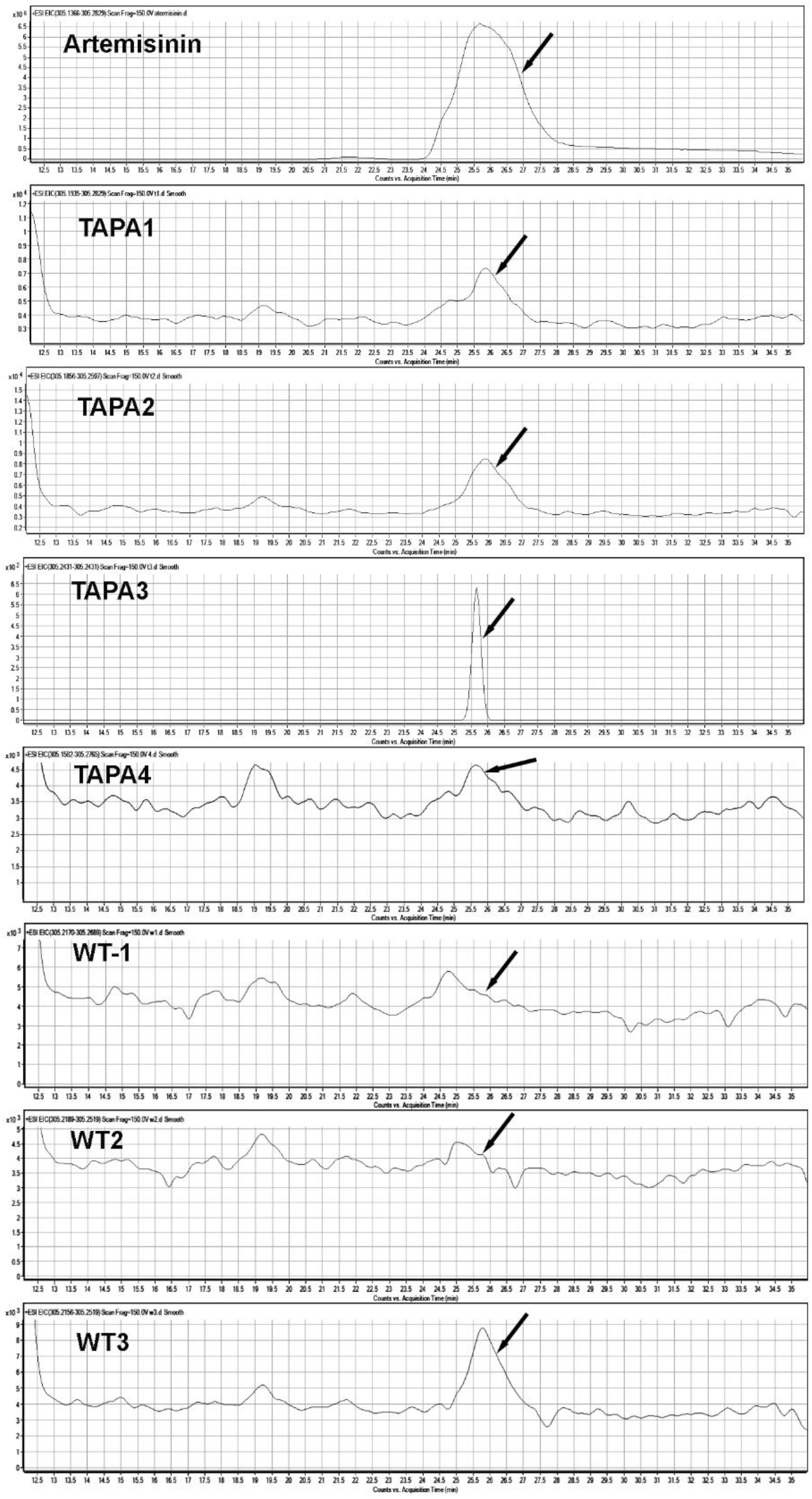
Elective ion chromatography of artemisinin. Authentic artemisinin was used as standard. Artemisinin was detected in extracts of four TAPA1 cell types and three wild type cells. Arrows indicate artemisinin peaks.

**Fig. S7.**
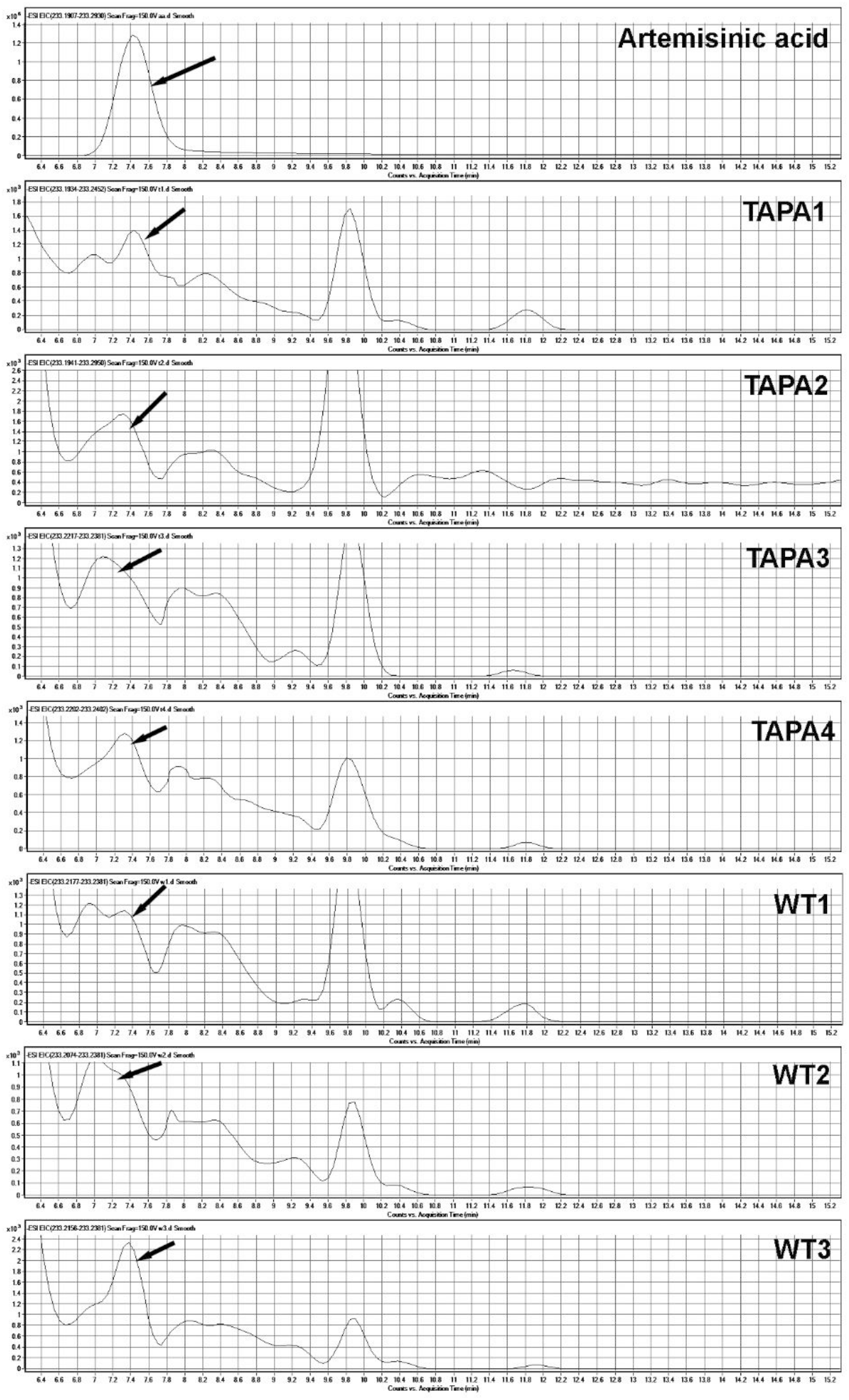
Elective ion chromatography of artemisinic acid. Authentic artemisinic acid was used ad standard. Artemisinic acid was detected in extracts of four TAPA1 cell types and three wild type cells. Arrows indicate artemisinic acid peaks.

**Fig. S8.**
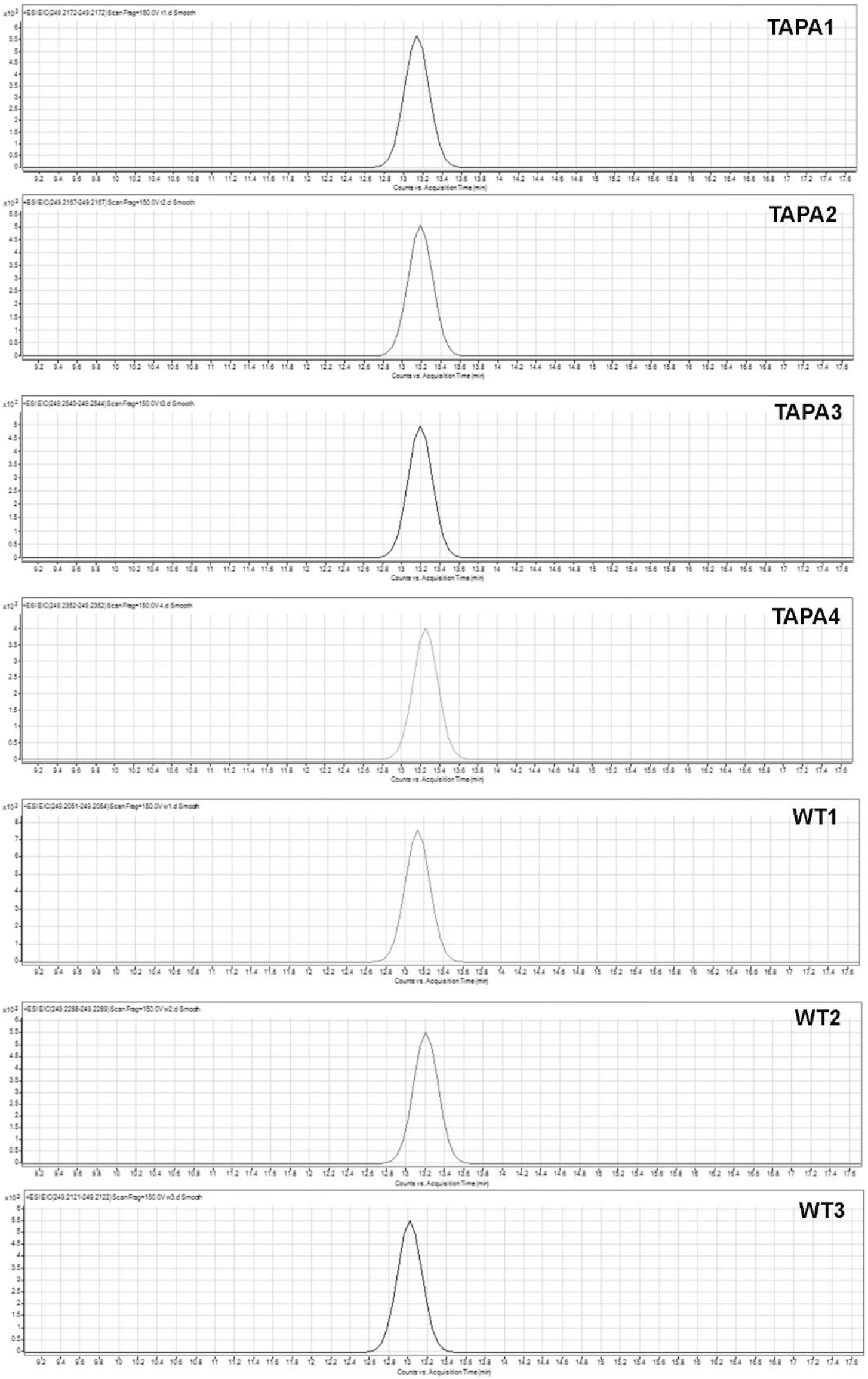
Elective ion chromatography of arteannuin b. Arteannuin b was detected in extracts of four TAPA1 cell types and three wild type cells. Arrows indicate arteannuin b peaks.

**S-Table 1.**
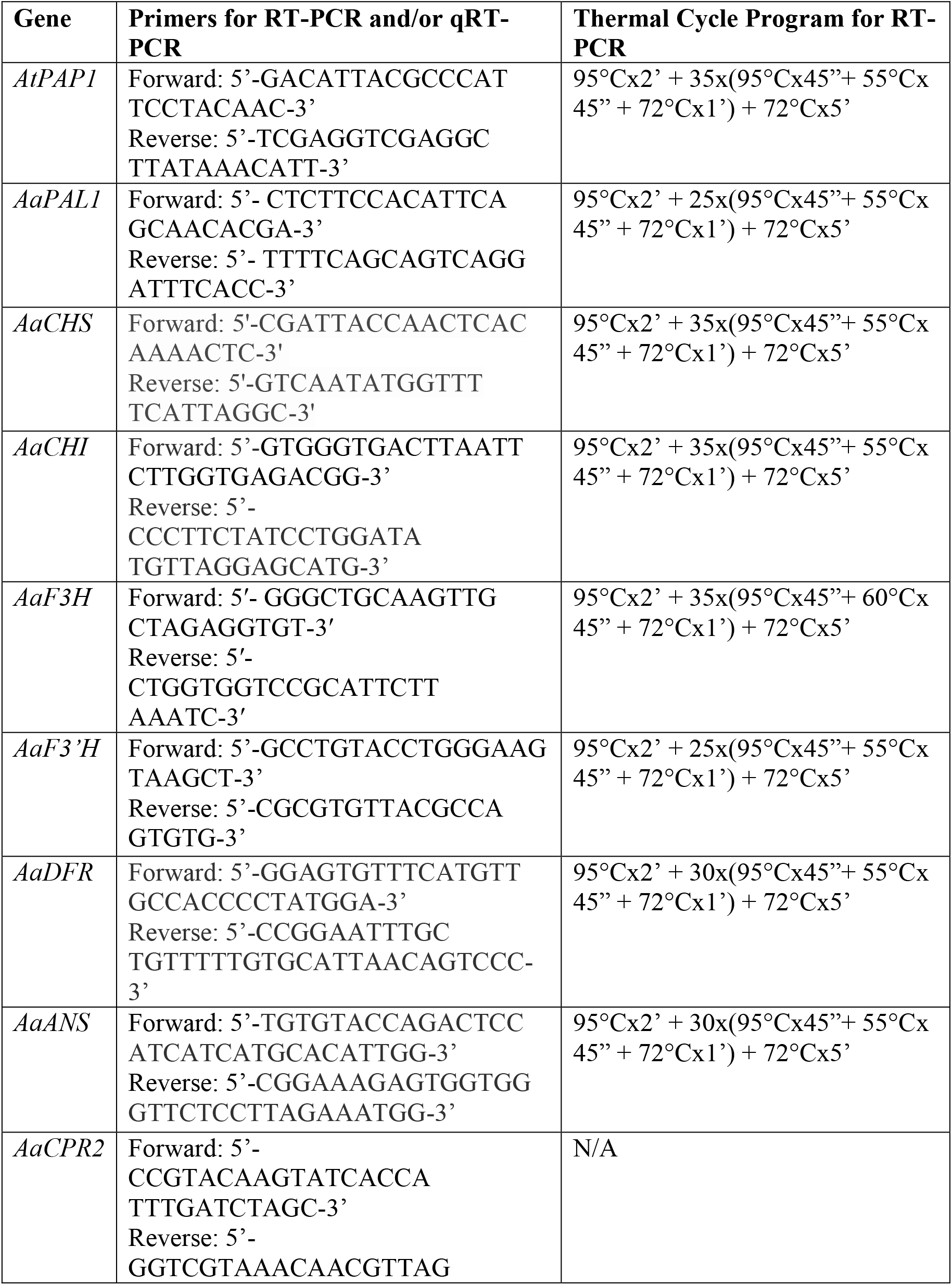
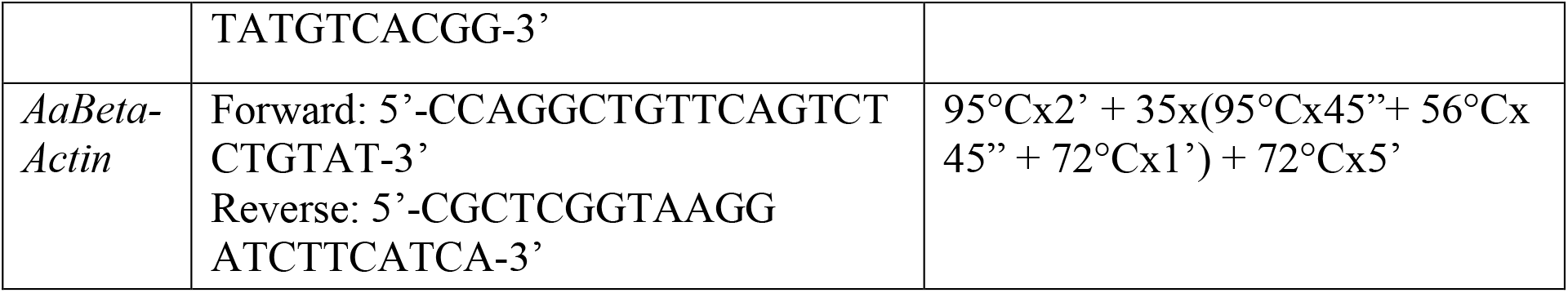
Primers for Semi-quantitative RT-PCR and qRT-PCR.

